# Evaluation of Tetraspanins in Extracellular Vesicle Bioengineering

**DOI:** 10.64898/2026.01.13.699196

**Authors:** Doste R. Mamand, Oskar Gustafsson, Helena Sork, Rim Jawad Wiklander, Safa Bazaz, Xiuming Liang, Vicky W.Q. Hou, Dhanu Gupta, André Görgens, Joel Z. Nordin, Samir EL Andaloussi, Oscar P.B Wiklander

**Affiliations:** Department of Laboratory Medicine, Division of Biomolecular and Cellular Medicine, Karolinska Institutet, Stockholm, Sweden; Breast Center, Karolinska Comprehensive Cancer Center, Karolinska University Hospital, Stockholm, Sweden; Biology Research center, Research Center, University of Zakho, Zakho, 42002, Duhok, Kurdistan Region, Iraq; Institute of Technology, University of Tartu, Tartu, Estonia; Karolinska ATMP Center, ANA Futura, Huddinge, Sweden; Department of Cellular Therapy and Allogeneic Stem Cell Transplantation (CAST), Karolinska University Hospital, Huddinge, Sweden; Center for Hematology and Regenerative Medicine (HERM), Department of Medicine Huddinge, Karolinska Institutet, SE-141 83 Stockholm, Sweden; Center of Research and Strategic Studies, Lebanese French University, Erbil 44001, Iraq; Department of Clinical Immunology and Transfusion Medicine (KITM), Karolinska University Hospital, Stockholm, Sweden; Department of Paediatrics, University of Oxford, Oxford, UK

**Author notes:** These authors contributed equally (Oskar Gustafsson, Helena Sork). These authors contributed equally (Joel Z. Nordin, Samir EL Andaloussi, Oscar P.B Wiklander).

**Keywords:** CD9, CD63, CD81, CRISPR-Cas9, EV release, EVs Bioengineering

## Abstract

Extracellular vesicles (EVs) are nano-scale structures produced by cells that transport biological substances for intercellular communication. The tetraspanins CD9, CD81, and CD63 are crucial to EV biogenesis and function. This study uses CRISPR-Cas9 system to knock out (KO) CD9, CD63, and CD81 in HEK293T cells. The goal is to investigate the role of these tetraspanins in EV bioengineering with the hypothesis that repressing endogenous production may increase the availability of exogenously introduced tetraspanin-fusion constructs and increase engineered EV production.

Firstly, it is observed that individually knocking out a tetraspanin does not significantly affect EV formation. However, when all three tetraspanins are simultaneously knocked out, there is a marked decrease in EV production, as measured by nanoparticle tracking analysis (NTA). Secondly, upon reintroduction of the corresponding tetraspanins fused to firefly ThermoLuc (Tluc) or neon green (mNG) into the PanKO-, CD9KO, CD63KO-, and CD81KO-cells, the engineered EVs display a significant increase in production by 50% to 70% compared to transduction of wild-type (WT) cells, as measured by luminometer and imaging flow cytometry.

These findings emphasize the potential of tetraspanin KO in the bioengineering of EVs, paving the way for new therapeutic applications by enhancing production and potentially modifying their cargo.

## 1. Introduction

Extracellular vesicles (EVs) are critical mediators of intercellular communication and have become a central focus in modern biomedical research due to their functional versatility and clinical potential[1] These nanoscale vesicles are released by a broad spectrum of cell types and are detectable in various biological fluids, including blood, urine, and saliva[2] EVs encapsulate and transport a wide range of bioactive molecules such as proteins, lipids, and RNA species that enable them to influence both physiological and pathological processes in recipient cells[3] Based on their size, biogenesis, and mechanisms of release, EVs are generally categorized into three major subtypes. Exosomes (30–150 nm) originate from the endosomal system via inward budding of the endosomal membrane, resulting in the formation of intraluminal vesicles within multivesicular bodies, which are subsequently released upon fusion with the plasma membrane. In contrast, microvesicles (100–1000 nm) are generated through direct outward budding from the plasma membrane, while apoptotic bodies (500–2000 nm) are produced during the execution phase of apoptosis and contain fragments of cellular components[4] EVs have been implicated in a range of physiological processes[5], including immunomodulation[6] tissue repair and regeneration[7]and intracellular signaling[8]They are also involved in various pathological conditions, such as hematopoietic dysregulation[9], tumor progression[10], and neurodegenerative disorders[11] Due to their inherent stability and ability to preserve and transport molecular cargo over long distances, EVs have emerged as promising tools for both therapeutic delivery and diagnostic applications[12], [13], [14], [15] and are increasingly regarded as valuable biomarkers and targets for clinical intervention[14].

A hallmark feature of EVs is their enrichment in tetraspanins, a superfamily of transmembrane proteins known for organizing specialized membrane domains and coordinating diverse molecular interactions. Tetraspanins interact with a broad array of transmembrane and cytosolic signaling partners, contributing to the assembly of tetraspanin-enriched microdomains (TEMs) that play critical roles in cellular signaling and membrane organization[16], [17]. Structurally, tetraspanins are characterized by four transmembrane domains, conserved extracellular cysteine residues, and specific polar amino acids within their transmembrane segments. Most members also contain palmitoylation sites and are frequently glycosylated, which further influences their functional properties[18]. These proteins participate in numerous essential cellular processes, including adhesion, motility, invasion, membrane fusion, intracellular signaling, and protein trafficking[19]. The functional specificity of tetraspanins is largely governed by the spatial distribution and organization of five key molecular components within the membrane microdomains, which collectively regulate membrane dynamics, trafficking, and signaling cascades[20], [21]. Among the commonly expressed tetraspanins are CD9, CD63, CD81, CD82, and CD151, while others such as Tssc6, CD37, and CD53 exhibit restricted expression patterns, particularly within hematopoietic lineages[22]. Immunoelectron microscopy has confirmed the localization of tetraspanins to endocytic compartments, supporting their widespread application as canonical markers for EV identification and characterization[23].

Within EV producing cells, CD9 is predominantly localized to the plasma membrane, CD63 is mainly confined to endosomal compartments, and CD81 is distributed across both subcellular sites[24]. However, the expression levels and subcellular localization of these tetraspanins are known to vary among different cell types. This variability is consistent with findings that CD81 and CD9 are frequently enriched in isolated EV populations, suggesting that some cell types may preferentially secrete ectosomes rather than exosomes[25]. Furthermore, recent studies have demonstrated that C-terminal modifications of CD63 disrupt its endosomal targeting signal, resulting in re-localization to the plasma membrane. This re-localization is associated with a more than sixfold increase in the secretion of CD63-positive EVs[26]. In previous investigations, CD9, CD63, and CD81 have been successfully employed to facilitate the incorporation of protein cargo into or onto EVs, thereby enabling the production of bioengineered vesicles for therapeutic and experimental applications[27], [28].

Optimizing the yield of EVs has become a critical priority in the field, particularly to facilitate the scalable production of EVs for therapeutic applications. A range of methodological approaches has been developed to enhance EV output in vitro systems. In this study, we aimed to investigate the role of tetraspanins, specifically CD9, CD63, and CD81, in the production and characteristics of EVs by generating HEK293T single-cell KO models using CRISPR/Cas9 technology. By knocking out these tetraspanins individually and in combination, we assessed how their absence affected the yield of EVs. We also explored using these tetraspanins in engineering cells to produce EVs with specific surface proteins or luminal cargo. We constructed plasmids that fused the tetraspanins to ThermoLuc (Tluc) to achieve this[27]. It was hypothesized that introducing a fusion construct could lead to competition between endogenous tetraspanins (e.g., CD9, CD63, or CD81) and their corresponding fused versions (e.g., TLuc-CD9, TLuc-CD63, or TLuc-CD81). This competition could reduce the expression of the fusion constructs in WT cells.

The results showed that knocking out single tetraspanins did not significantly affect the production of EVs. However, knocking out all three tetraspanins resulted in a marked decrease in EV secretion. Our findings also revealed that cells lacking these specific tetraspanins did not face competitive inhibition. Instead, when engineered, these PanKO-, CD9KO, CD63KO-, and CD81KO-cells exhibited an increased production of engineered EVs and contained more of the desired cargo. This indicates that the absence of tetraspanins allows more efficient incorporation of engineered proteins into EVs. Our study provides valuable insights into the role of tetraspanin expression in EV bioengineering. Overall, it emphasizes the importance of tetraspanins in EV production and highlights the potential of using KO models to optimize engineered EV systems for therapeutic applications.

## 2. Results

### 2.1 Characterization of EVs and Cells

First, we aimed to determine whether KO of CD9, CD63, or CD81 affects EV release. To confirm the successful generation of single-cell KO models for these tetraspanins, we stained both the cells and their corresponding EVs with APC-conjugated antibodies. Tetraspanin expression was validated using flow cytometry and imaging flow cytometry for single-vesicle detection. Flow cytometric analysis showed that HEK293T WT cells and WT-EVs were positive for CD9, CD63, and CD81 on their surface (Figure 1B and 1C). In contrast, PanKO cells (lacking the tetraspanins CD9, CD63, and CD81) showed no detectable staining for CD9, CD63, or CD81, confirming the knockout efficiency; this absence of tetraspanin expression was also observed in PanKO-derived EVs, which lacked detectable levels of these markers (Figure 1B and 1C, S1D). Further analysis of single tetraspanin KO confirmed absence of respective tetraspanin of the cells and derived EVs with CD9KO displaying negative staining of CD9, CD63KO cells and EVs were negative for CD63, and CD81KO cells and EVs were negative for CD81 (Figure 1B and 1C). Overall, these findings validate the generation of tetraspanin KO cell models and demonstrate the successful elimination of tetraspanin expression in both cells and their corresponding EVs, setting the stage for further exploration of the functional roles of these proteins in EV biogenesis and bioengineering.

**Figure 1:**
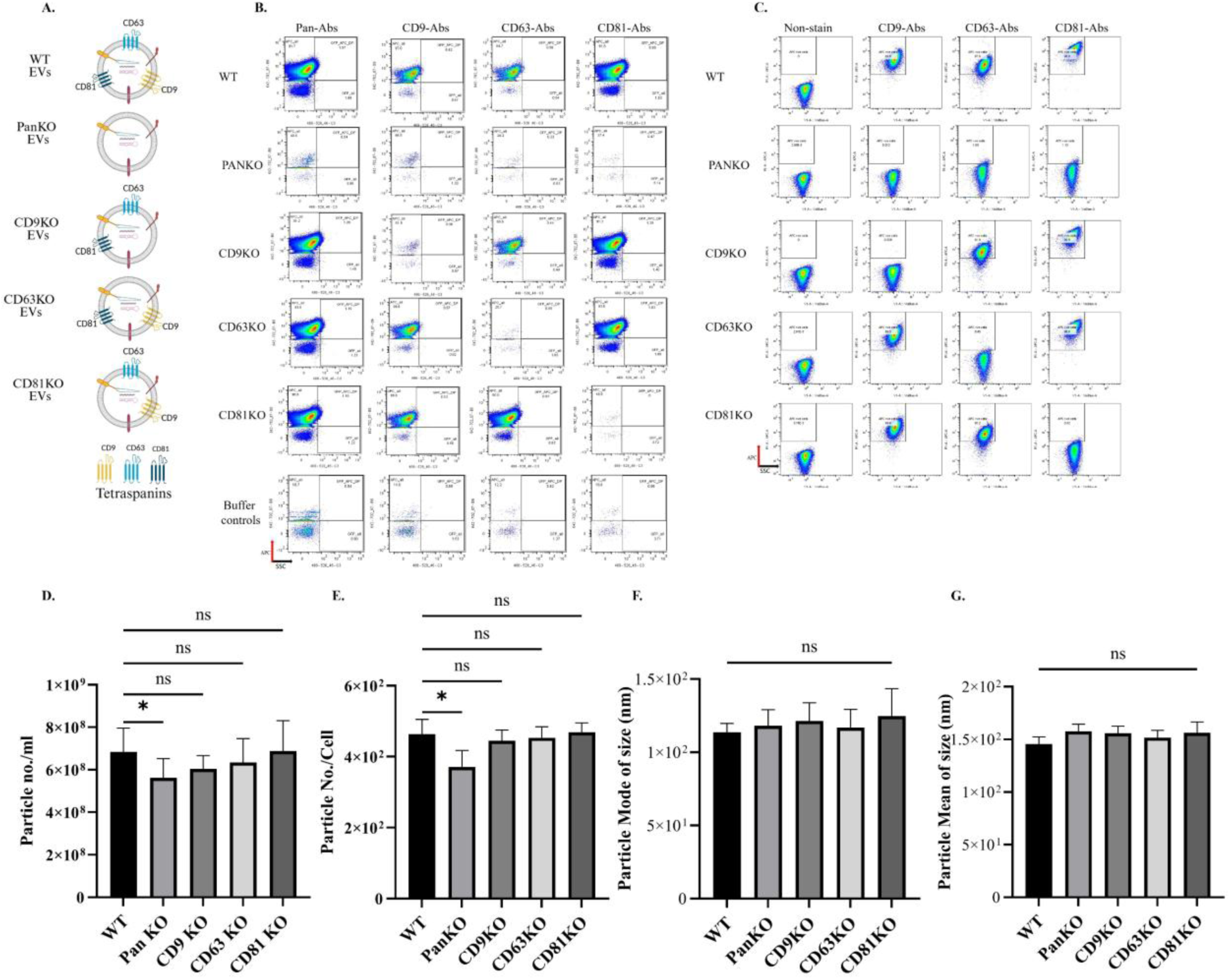
Flow cytometry validation of tetraspanins surface expression on the cell and the EVs after single-cell knockout of tetraspanins. A. Schematic illustration of tetraspanin distribution on EVs derived from individual tetraspanin KO cells: CD9KO, CD63KO-, and CD81KO-cells, PanKO-, and WT-cells (created with BioRender.com). B. Surface expression of EV-associated tetraspanins was assessed by single-staining with APC-conjugated pan-tetraspanin antibodies, followed by single EV imaging flow cytometry (analyzed with FlowJo v10.6.1). C. Cellular surface expression of tetraspanins was evaluated using APC-conjugated pan-tetraspanin antibodies and conventional flow cytometry (analyzed with FlowJo v10.6.1). D–G. NTA was used to quantify and characterize EVs. D. Total number of EVs produced per milliliter by each cell line. E. Particle count normalized to cell number. F. Mode size of EVs, represented by diameter in nanometers. G. Mean size of EVs, represented by diameter in nanometers. The data are presented as means (±SD, n = 6). Statistical analysis was performed using one-way ANOVA; significance is indicated as *p < 0.05.

Next, we conducted a WST-1 assay to evaluate the effects of tetraspanin KO on cellular proliferation. The aim was to assess whether the KO of these tetraspanins affected cell growth. There were no significant differences in proliferation degree between the KO and WT cells (Figure S1A). Cell viability and concentrations were also assessed using trypan blue exclusion and were comparable across all groups (Figures S1B and S1C). Moreover, the overall protein concentrations remained consistent among all cell lines, as illustrated in Figure S1E. This consistency indicates that the tetraspanin KO did not adversely modify the general characteristics of the cells. To further investigate the functionality of the PanKO-, CD9KO, CD63KO-, and CD81KO-cells in terms of transfection efficiency, we transfected the cells using eGFP plasmids linked to CD9, CD63, and CD81. The results showed that the transfection efficiency remained stable across the KO cell lines when using Lipofectamine 2000, as depicted in Figure S1F and S1G. It indicates that the KO of tetraspanins did not impair the ability of these cells to take up and express plasmid DNA effectively. These findings suggest that the KO of tetraspanins does not significantly impact cell proliferation, viability, or transfection efficiency, allowing for further investigation into their roles in EV bioengineering.

### 2.2 Quantification of EVs produced by KO and WT cells

Next, we investigated the roles of CD9, CD63, and CD81 on EV yield from the different KO cell lines. First, the EVs were characterized by NTA and TEM (Figure S2 and S3).The NTA results indicated no statistically significant differences in EV production between single tetraspanin PanKO-, CD9KO, CD63KO-, and CD81KO-cells and WT cells; however, the overall quantity of EVs produced by the single KO cells: CD9KO, CD63KO-, and CD81KO-cells was lower than that of WT cells. Notably, CD9KO showed slightly lower EV production than CD63KO and CD81KO cells, as illustrated in Figure 1D and 1E. Statistically significant differences in EV production were observed between PanKO cells and WT cells, indicating that the combined KO of all three tetraspanins had a more pronounced effect on EV production. When comparing the single tetraspanin: CD9KO, CD63KO-, and CD81KO-cells to the PanKO cells, no significant variations in EV release were found between CD9KO-, CD63KO-, and CD81-KO cells. Importantly, despite the variations in EV production, no difference regarding measured diameter values obtained by NTA were detected across all cell lines, regardless of the presence or absence of CD9, CD63, or CD81 (Figure 1F-Gand Figure S2). These findings suggest that while the combined KO of all three tetraspanins significantly affects EV release, the impact of individual tetraspanin CD9, CD63, or CD81 KO is relatively modest. This study underscores the complexity of tetraspanin roles in EV production and highlights the potential for utilizing single tetraspanin KO cells: CD9KO-, CD63KO-, and CD81KO-cells in future EV engineering applications.

### 2.3 Generation of stable luminescent protein expression in KO and WT cells by introducing Tluc-CD9, Tluc-CD63, and Tluc-CD81

Next, we tested the hypothesis that competition between endogenous and exogenous tetraspanins interferes with efficient EV engineering. To investigate this, we generated stable luminescent expression in both WT and PanKO-, CD9KO, CD63KO-, and CD81KO-cells by introducing Tluc-tagged CD9, CD63, and CD81 (TlucCD9, TlucCD63, and TlucCD81). This approach enabled functional studies of these tetraspanins through luminescence-based tracking.

To achieve stable expression, lentiviral vectors were employed, and Cerulean was co-expressed as an internal control (Supplementary Table 1). Notably, Cerulean was not fused to any EV sorting domains, thereby limiting its presence to the cellular compartment and preventing its incorporation into EVs (Figure S4). We used mean fluorescence intensity (MFI) as a metric to facilitate cell sorting based on Cerulean expression. Before sorting, cells were treated with 1000 µg/ml Zeocin to select successfully transduced cells (Figures 2B-D, 3B-D, and 4B-D). The expression of Cerulean in each cell line was verified using flow cytometry, confirming the successful integration and expression of the fluorescent marker. All cell lines were assessed for cellular proliferation to determine whether the transduction process affected their proliferation (Figure S7, S10, S13).

**Figure 2:**
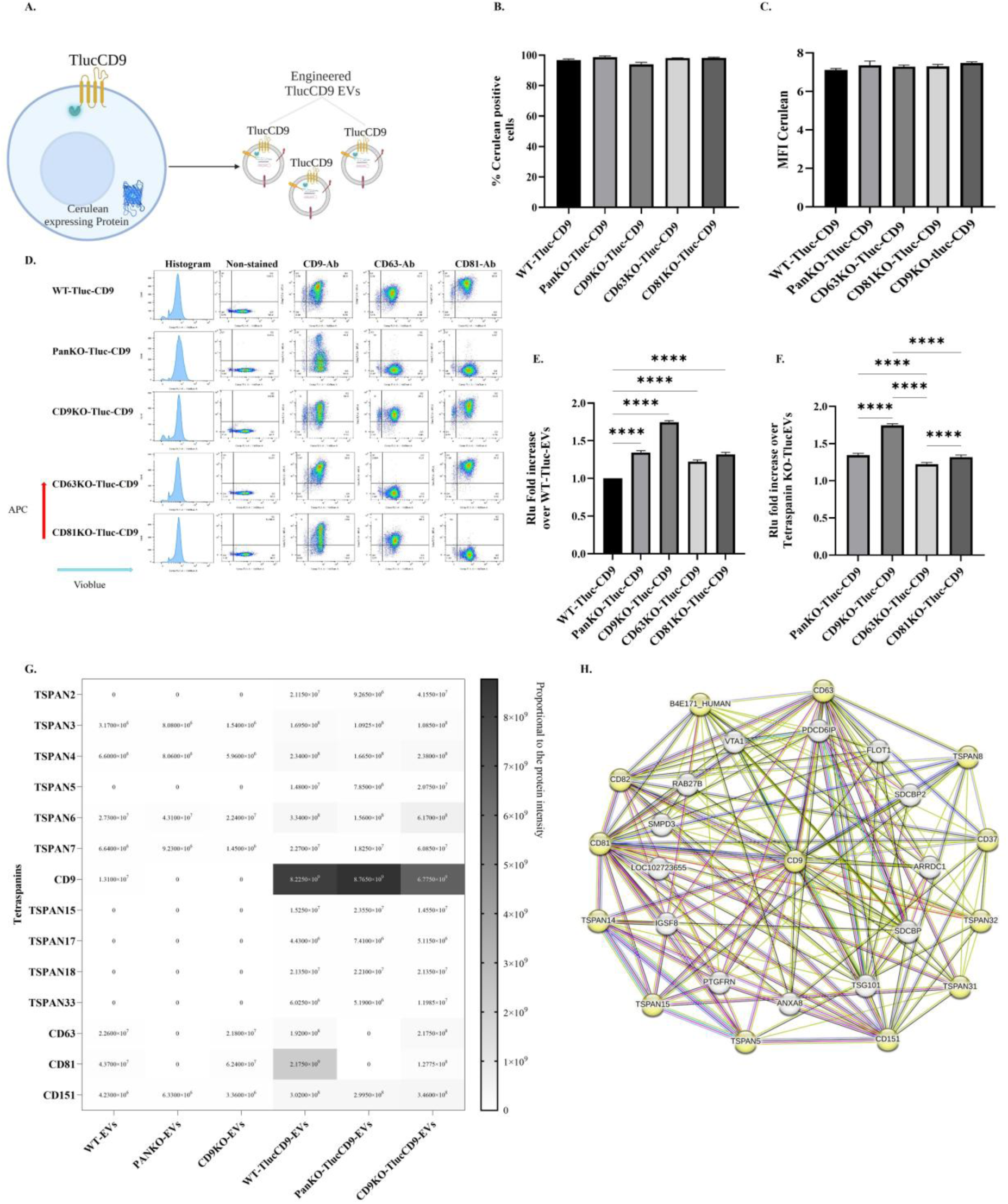
Lentiviral transduction to generate WT-, PanKO-, CD9KO-, CD63KO-, and CD81KO-cells expressing TlucCD9-Cerulean. A. Schematic workflow of generating engineered Tluc-EVs by introducing TlucCD9-Cerulean lentiviruses (created with BioRender.com). B. The percentage of Cerulean positive cells. C. The mean fluorescence intensity (MFI) of Cerulean positive cells. D. Flow cytometry plot for the cells after staining with APC-conjugated CD9/CD63/CD81 tetraspanin antibodies. E. Fold increase in engineered Tluc-CD9 EVs in PanKO-, CD9KO, CD63KO-, and CD81KO-cells over WT cells (Data are normalized to the RLU of EVs from WT cells). F. Fold increase of engineered Tluc-CD9EVs over PanKO-, CD9KO, CD63KO-, and CD81KO-cells (Data are normalized to the RLU of EVs from WT cells and compared across KO groups). G. Heatmap of tetraspanins in EVs. H. Interaction network of CD9 with the tetraspanins retrieved from STRING. The data are presented as means (±SD, n = 3-5). One-way ANOVA was used to show significance and was illustrated as follows: **** p < 0.0001.

**Figure 3:**
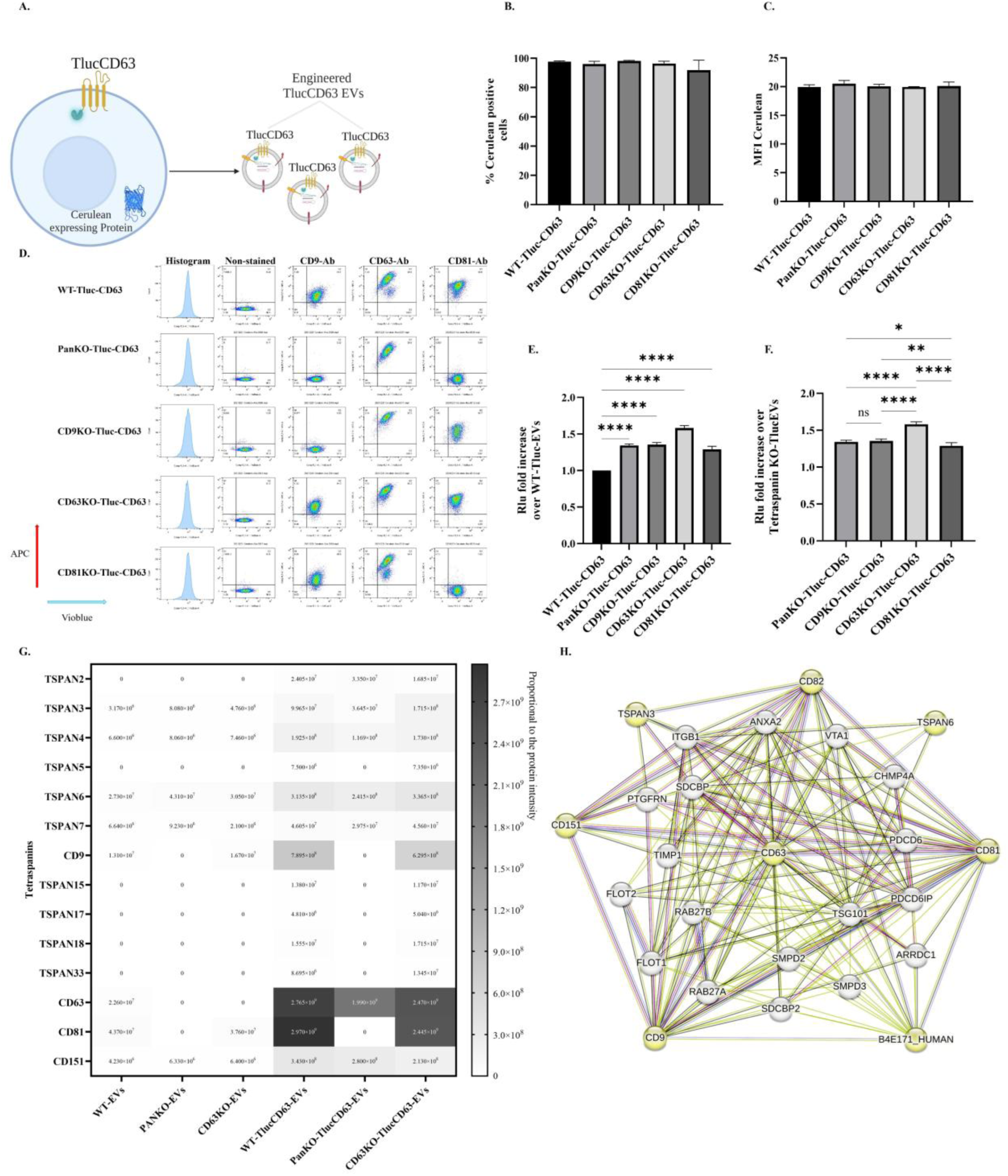
Lentiviral transduction to generate WT, PanKO, CD9KO, CD63KO, and CD81KO cells expressing TlucCD63-Cerulean. A. Schematic workflow of engineered Tluc-EVs by introducing TlucCD63-Cerulean lentiviruses (created with BioRender.com). B. The percentage of Cerulean positive cells. C. The mean fluorescent intensity (MFI) of Cerulean positive cells. D. Flow cytometry plot for the cells after staining with APC-conjugated CD9/CD63/CD81 tetraspanin antibodies. E. Fold increase of engineered Tluc-CD63 EVs in PanKO-, CD9KO, CD63KO-, and CD81KO-cells over WT cells (Data are normalized to the RLU of EVs from WT cells). F. Fold increase of engineered Tluc-CD63EVs in between PanKO-, CD9KO, CD63KO-, and CD81KO-cells (Data are normalized to the RLU of EVs from WT cells and compared across KO groups). G. Heatmap of expressing tetraspanins in EVs. H. Interaction network of CD63 with the tetraspanins retrieved from STRING. The data are presented as means (±SD, n = 3-5). One-way ANOVA was used to show significance and was illustrated as follows: * p < 0.05, ** p < 0.01, **** p < 0.0001.

**Figure 4:**
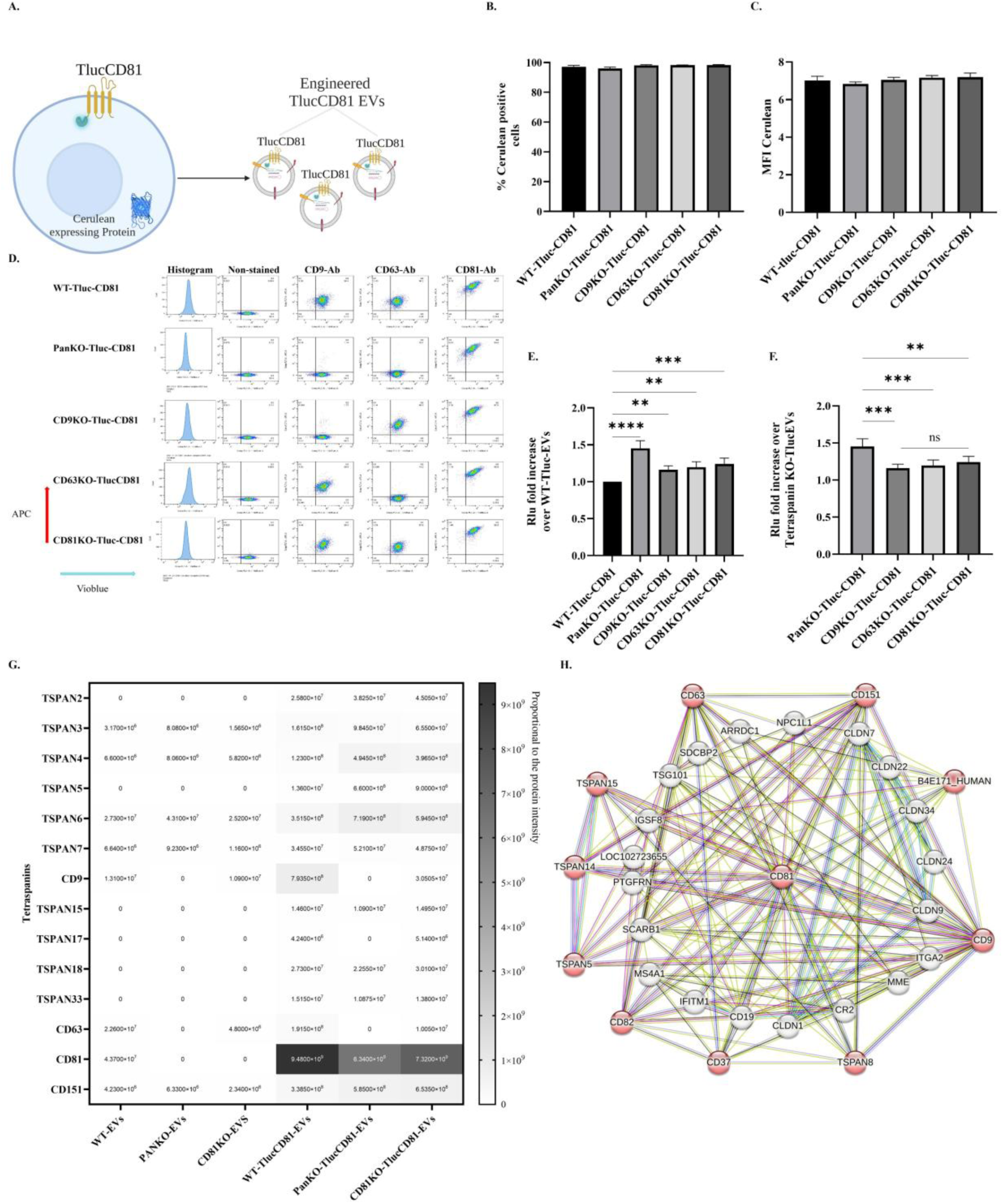
Lentiviral transduction to generate WT-, PanKO-, CD9KO-, CD63KO-, and CD81KO-cells expressing TlucCD9-Cerulean. A. Schematic workflow of engineered Tluc-EVs by introducing TlucCD81-Cerulean lentiviruses (created with BioRender.com). B. The percentage of Cerulean positive cells. C. The MFI of Cerulean positive cells. D. Flow cytometry plot for the cells after staining with APC-conjugated CD9/CD63/CD81 tetraspanin antibodies. E. Fold increase of engineered Tluc-CD81 EVs in PanKO-, CD9KO, CD63KO-, and CD81KO-cells over WT cells (Data are normalized to the RLU of EVs from WT cells). F. Fold increase of engineered Tluc-CD81EVs in between PanKO-, CD9KO, CD63KO-, and CD81KO-cells (Data are normalized to the RLU of EVs from WT cells and compared across KO groups). G. Heatmap of expressing tetraspanins in EVs. H. Interaction network of CD81 with the tetraspanins retrieved from STRING. The data are presented as means (±SD, n = 3-5). One-way ANOVA was used to show significance and was illustrated as follows: ** p < 0.01,** * p < 0.001,**** p < 0.0001.

For subsequent experiments, we used the same MFI of Cerulean as a standard internal control value across all cell lines, ensuring comparison consistency. This approach allowed us to assess the effect of the reintroduced Tluc-CD9, -CD63 and -CD81 on EV engineering and characteristics while controlling for any potential variability in cell viability or transduction efficiency. By employing this strategy, we aimed to elucidate how the presence of exogenous tetraspanins might influence the yield of engineered Tluc-EVs in both KO and WT backgrounds, contributing valuable insights into EV bioengineering.

### 2.4 Introducing Tluc-CD9, Tluc-CD63, and Tluc-CD81: Enhancing the level of engineered Tluc-EVs in PanKO-, CD9KO, CD63KO-, and CD81KO-cells

To assess whether PanKO-, CD9KO, CD63KO-, and CD81KO-cells could produce more engineered Tluc-EVs than WT cells when expressing the TlucCD9 (Figure 2A), EVs from cell lines (with matching MFI of Cerulean) were purified from the CM and quantified and characterized using NTA and TEM (Figure S5, S6, S7B-C). For relative luminescent unit (RLU) of Tluc, we used an equal number of EVs from each cell line based on NTA measurements.

The results indicated that reintroducing CD9 tagged to Tluc significantly increased engineered Tluc-EV production in all PanKO-, CD9KO, CD63KO-, and CD81KO-cells by 30-70% compared to WT cells (Figure 2E). CD9KO-TlucCD9 cells produced 50% more engineered Tluc-EVs than PanKO-TlucCD9, CD63KO-TlucCD9, and CD81KO-TlucCD9 cells (Figure 2F). Interestingly, while PanKO-TlucCD9 cells, lacking endogenous CD9, did not demonstrate the same increase, they still produced 20% more engineered Tluc-EVs than CD63KO-TlucCD9 cells, although not as much as CD81KO-TlucCD9 cells (Figure 2F and Figure S7D).

PanKO-, CD9KO, CD63KO-, and CD81KO-cells exhibited an overall increase in the release of engineered Tluc-EVs, with CD9KO and PanKO cells showing the most pronounced enhancement following reintroduction of Tluc-CD9. This observation prompted an investigation into the underlying mechanisms. Mass spectrometry-based proteomic analysis revealed the presence of multiple tetraspanins beyond the commonly used in EV research CD9, CD63, and CD81, including TSPAN2, TSPAN5, TSPAN15, TSPAN17, TSPAN18, and

TSPAN33 (Figure 2G and Figure S7E-F). Interestingly, the tetraspanin expression profile differed depending on the specific KO cell line. In addition to the loss of targeted tetraspanins, altered expressions were observed for TSPAN3, TSPAN4, TSPAN6, TSPAN7, and TSPAN151, which may influence EV composition and potential engineering strategies. Furthermore, STRING-based predictions revealed interactions between CD9 and multiple other tetraspanins (including TSPAN5, TSPAN8, TSPAN14, TSPAN15, TSPAN31, and TSPAN32), as well as with CD82, CD37, B4E171, CD151, CD63, and CD81 (Figure 2H). These findings suggest that CD9 may compete with other tetraspanins and EV-sorting proteins (such as ARRDC1 and PTGFRN) when reintroduced, potentially influencing the yield and composition of engineered Tluc-EVs.

We next investigated the effects of reintroducing TlucCD63 on engineered EV production in both KO and WT cells (Figure 3A). After quantification and characterization of the EVs using NTA and TEM (Figure S8, S9, S10B-C). We observed, using a similar experimental approach, that all PanKO-, CD9KO, CD63KO-, and CD81KO-cells expressing TlucCD63 produced engineered Tluc-EVs at levels 40% to 70% higher than their WT counterparts (Figure 3E), consistent with previous findings for TlucCD9. Notably, among the different KO cell lines, CD63KO-TlucCD63 cells exhibited a marked increase in Tluc-EV production approximately 40% to 50% more compared to PanKO-TlucCD63, CD9KO-TlucCD63, and CD81KO-TlucCD63 cells (Figure 3F and Figure S5D). This pattern parallels the enhanced EV production observed in CD9KO-TlucCD9 cells. However, similar to cells transduced with TlucCD9, we found that multiple tetraspanins were endogenously expressed and even overexpressed following TlucCD63 transduction (Figure 3G and Figure S5E–F). This suggests potential competition between exogenously expressed TlucCD63 and endogenous tetraspanins, which may limit the overall formation of engineered Tluc-EVs. Comparable to TlucCD9, STRING analysis showed strong interactions between CD63 and several other tetraspanins and non-tetraspanins (Figure 3H).

Next, we assessed the impact of reintroducing TlucCD81 on engineered EV production (Figure 4A). After quantification and characterization of the EVs using NTA and TEM (Figure S11, S12, S13B-C). Employing the same experimental strategy, we observed that all KO cell lines expressing TlucCD81 produced 30% to 50% more engineered Tluc-EVs compared to WT cells (Figure 4E). Importantly, the MFI and the percentage of Cerulean-positive cells used as an internal control remained consistent across all cell lines, confirming comparable transduction efficiency and expression levels (Figure 4C–D).

Interestingly, unlike observations with TlucCD9 and TlucCD63, PanKO-TlucCD81 cells exhibited a marked increase in engineered EV production, generating approximately 40% more engineered Tluc-EVs compared to CD9KO-TlucCD81, CD63KO-TlucCD81, and CD81KO-TlucCD81 cells (Figure 4F and Figure S13E-F). In contrast, CD81KO-TlucCD81 cells did not show significant differences in Tluc-EV output relative to the other single KO groups, underscoring the complex and context-dependent nature of tetraspanin involvement in EV biogenesis.

Similar to the findings observed with TlucCD9 and TlucCD63, transduction with TlucCD81 led to the expression and overexpression of various other tetraspanins (Figure 4G and Figure S6B-C). Furthermore, STRING analysis demonstrated significant interactions between CD81 and multiple EV-sorting proteins (Figure 4H), underscoring a competitive and interdependent protein network that regulates engineered EV production and the necessity of comprehending these interactions.

### 2.5 Reintroducing CD63 and CD81, tagged to a green fluorescent protein, and stably expressing in KO and WT cells enhances the levels of engineered mNG-EVs

Next, to determine whether alternative strategies for engineered EV production in KO-versus WT-cells would yield results consistent with the luminescence assay, we transduced WT-cells and PanKO-, CD9KO, CD63KO-, and CD81KO-cells with either CD63-mNeonGreen (CD63-mNG) or CD81-mNG lentiviral constructs (Figure 5A and Figure S16A), enabling stable production of mNG-labeled EVs. Each cell line was seeded at 10 × 10⁶ cells, with comparable MFI and percentage of mNG-positive cells (Figure 5B–D and Figure S16B–D). Cells were cultured in 20 mL of either serum-free Opti-MEM or serum-supplemented DMEM for 48 hours.

**Figure 5:**
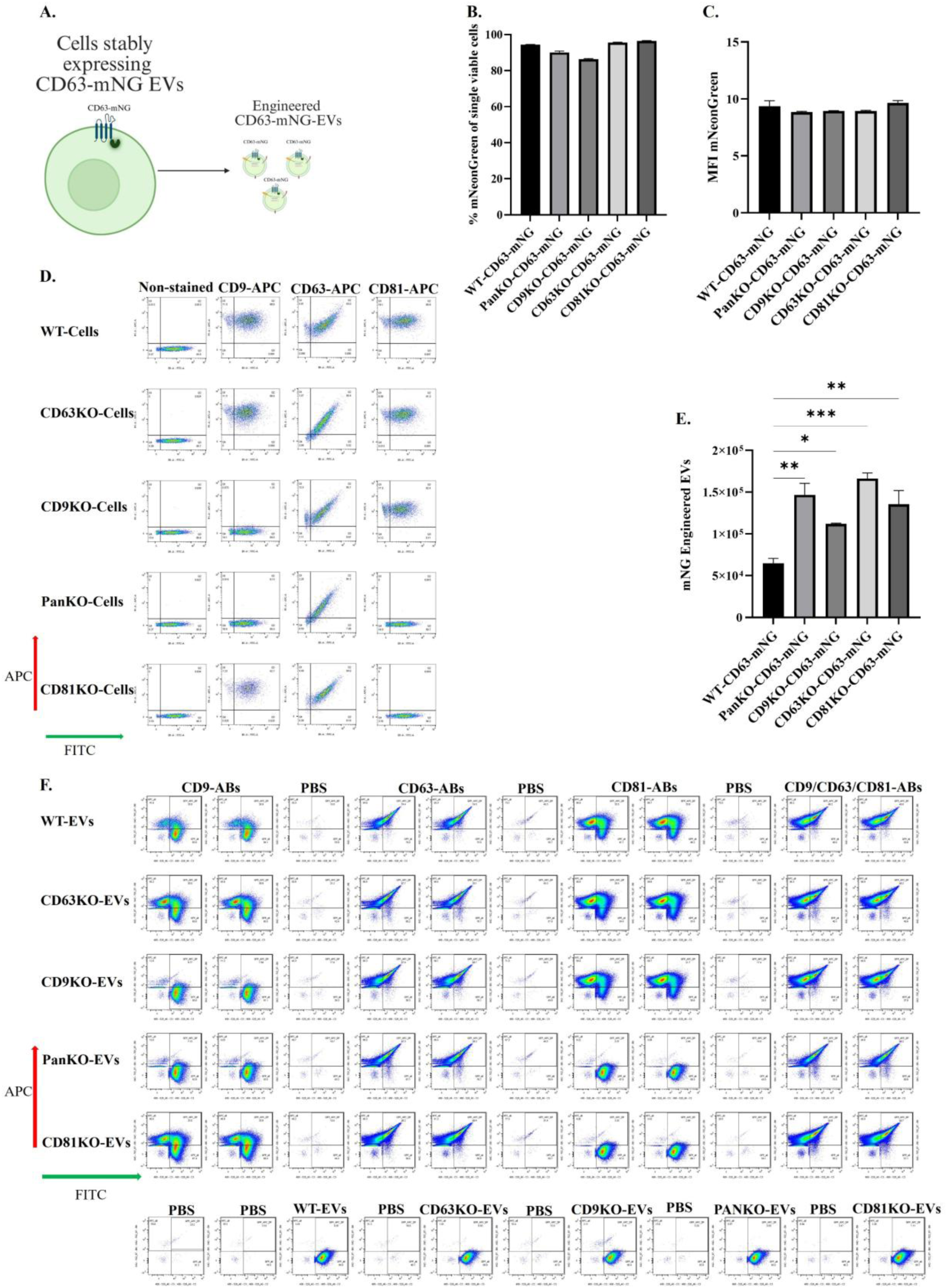
Generation of CD63-mNG-EVs in WT, PanKO-, CD9KO-, CD63KO-, and CD81KO-cells. A. Schematic workflow of engineered mNG-EVs by introducing CD63-mNG lentiviruses “created with BioRender.com”. B. Percentage of mNG positive cells after transduction using flow cytometry. C. MFI of the cells using flow cytometry. D. The flow cytometry plot for the cells after transduction, stained with APC-conjugated CD9/CD63/CD81 tetraspanin antibodies. E. Quantification of engineered CD63-mNG EVs from 17 µL of CM collected from KO) and WT cells. F. Imaging flow cytometry plot for the mNG-EVs derived from stably expressing mNG cells. The data are presented as means (±SD, n = 2-3). One-way ANOVA was used to show significance and was illustrated as follows: * p< 0.05; ** p < 0.01; *** p < 0.001.

Following incubation, EVs were isolated and purified from the CM, and their concentrations were assessed via NTA (Figure S14 and S15). Quantification of engineered mNG-EVs was performed using imaging flow cytometry (Amnis CellStream, Luminex, USA), following established EV analysis protocols[29]. The results showed that PanKO-, CD9KO, CD63KO-, and CD81KO-cells released significantly more mNG-positive EVs compared to WT cells when transduced with CD63-mNG (Figure 5E and 5F). Notably, the percentage and MFI of mNG expression were consistent across all cell lines, confirming that differences in EV output were due to tetraspanin knockout rather than variation in transduction efficiency or cell viability (Figure 5C).

To further validate these findings, we examined whether PanKO-, CD9KO, CD63KO-, and CD81KO-cells producing mNG-EVs continue to exhibit enhanced EV release in serum-containing conditions, as previously observed with engineered Tluc-EVs (Figure S16A). WT cells and PanKO-, CD9KO, CD63KO-, and CD81KO-cells were seeded at 10 × 10⁶ cells in DMEM supplemented with 10% FBS and transduced with CD81-mNG lentivirus. After 48 hours, EVs were isolated from the CM using a combination of spin filtration and SEC. Imaging flow cytometry analysis confirmed that PanKO-, CD9KO, CD63KO-, and CD81KO-cells released increased levels of mNG-positive EVs compared to WT cells (Figure S16E and S16F), supporting the conclusion that tetraspanin knockout enhances engineered EV production across different expression systems and culture conditions.

### 2.6 Reintroducing CD63 to CD63KO-cells restores and elevates CD63 levels in CD63KO-EVs

We next investigated whether CD63 expression could be restored to WT levels in a CD63KO background. For that we performed tandem mass spectrometry analysis of WT EVs, PanKO-, CD9KO-, CD63KO-, and CD81KO-EVs and EVs originating from TlucCD63 OE cells on either WT or CD63KO background. Our results showed that upon employment of the Tluc-CD63 construct, the level of CD63 can not only be restored, but even elevated beyond the WT background (Figure 6A). Specifically, upon reintroducing CD63 into KO cells, the level of CD63 in EVs, could be elevated more than 11-fold (log₂ fold change 11.19, adjusted p-value 8.27E-04, FDR 5%) as compared to more than 6-fold increase (log₂ fold change 6.15, adjusted p-value 0.0446, FDR 5%) obtained in WT EVs (Figure 6B).

**Figure 6:**
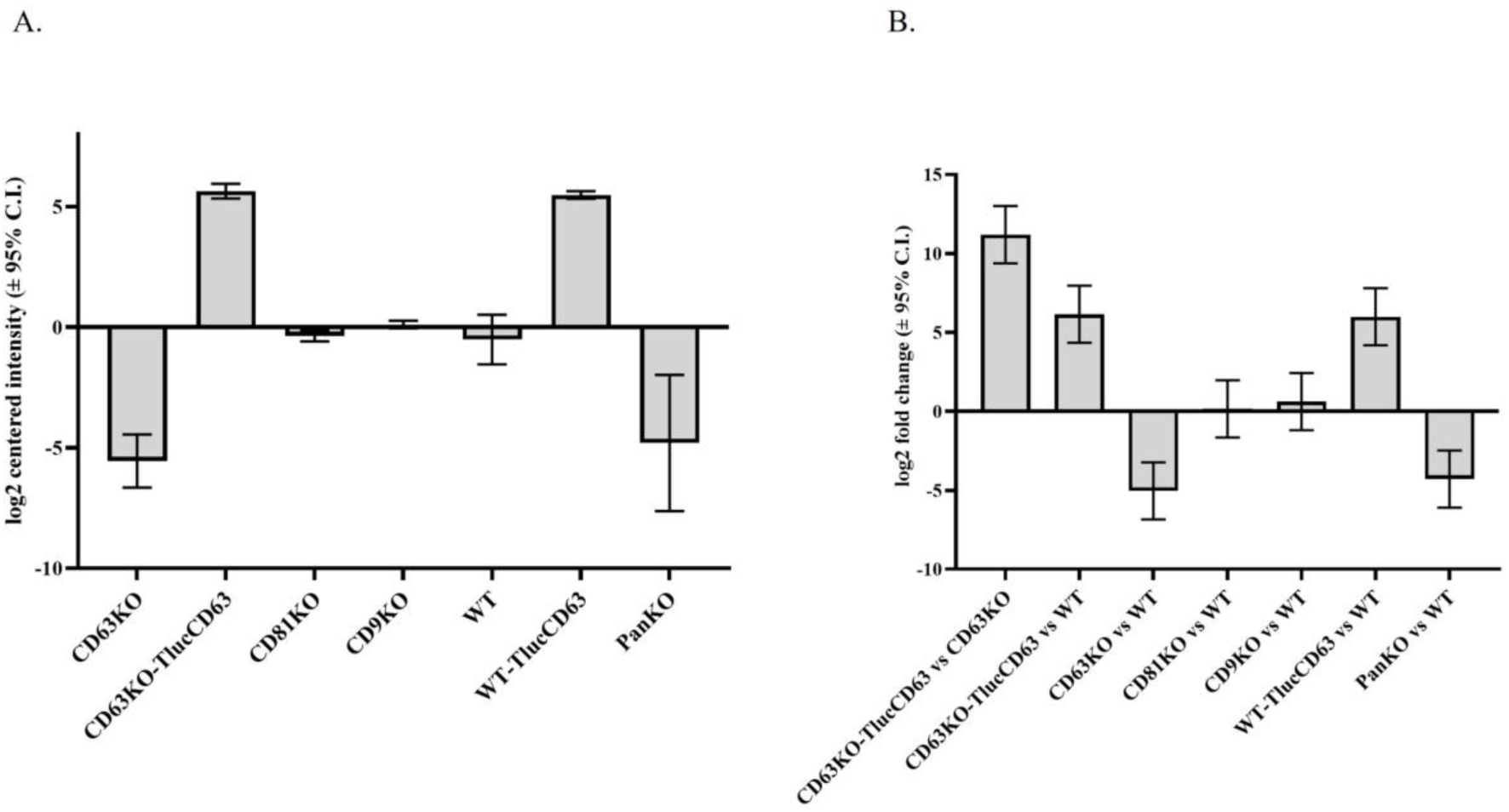
Proteomic evaluation on the restoration and expression of CD63 in EVs. (A) Expression of CD63 in EVs originating from CD63KO and WT HEK293T cells. The graph indicates the relative difference in CD63 level across samples using log2-transformed protein intensities centered across all samples. (B) Expression of CD63 in EVs from CD63KO cells transfected with Tluc-CD63 compared to CD63KO and WT cells. The graph indicates the log2 fold change in the relative CD63 level over CD63 KO EVs or WT EVs. Results represent data from three biological replicates.

## 3. Discussion

This study investigates the role of the tetraspanins CD9, CD63, and CD81 in EV release and bioengineering. Using the CRISPR/Cas9 system, we generated stable HEK293T KO cell lines targeting each tetraspanin individually (CD9, CD63, CD81) and in combination (PanKO). Our initial objective was to examine how these KOs affect EV secretion. Individual knockouts of CD9, CD63, or CD81 did not significantly impact EV release, consistent with prior studies [26]. However, it contrasts with previous findings where CD9 KO in SKMEL147 and CD81 KO in MDA-MB-231 cells increased EV release[30], [31], while KO CD9, CD81 or CD63 in MCF7 cells had no effect[32]. PanKO HEK cells showed a marked reduction in EV production, indicating that these tetraspanins act synergistically to support EV release. It is worth noting that the inconsistency among various studies challenges the notion of a general and critical role for tetraspanins in EV biogenesis. The variability across studies suggests that the influence of tetraspanins may be cell-type dependent or influenced by differences in EV isolation and analysis protocols. Despite the knockout of different tetraspanins, EV size remained consistent across all cell lines, as measured by NTA, suggesting that tetraspanins are not major regulators of EV size.

Secondly, we aimed to explore the effects of knocking out CD9, CD63, and CD81 on the bioengineering of EVs. We hypothesized that expressing fusion constructs of these tetraspanins would compete with their endogenous forms, potentially disrupting normal EV engineering processes. In contrast, this competition would be absent in tetraspanin KO HEK cells, thereby providing clearer insight into the individual roles these proteins play in EV bioengineering. While numerous studies have focused on improving the production and functionality of engineered Tluc-EVs for research and therapeutic applications, optimizing their yield remains a major challenge[33], [34]. Although the mechanisms for loading protein cargo often via tetraspanins such as CD9, CD63, and CD81 are relatively well characterized, efficient production of engineered Tluc-EVs is still difficult to achieve.

In our study, engineered Tluc-EVs were produced using stable luciferase-expressing cell lines, where a Tluc construct was fused to the N-terminus of CD9, CD63, or CD81. Cerulean protein served as an internal control, and engineered cells were sorted based on comparable Cerulean fluorescence intensity to ensure consistent MFI across cell lines. PanKO-, CD9KO, CD63KO-, and CD81KO-cells expressing Tluc-CD9, Tluc-CD63, or Tluc-CD81 exhibited increased production of engineered Tluc-EVs from 50% to 70% over WT cells when Tluc-CD9, Tluc-CD63, or Tluc-CD81 were overexpressed.

Notably, CD9KO-TlucCD9 cells showed a significant increase in engineered EV output compared to CD63KO-TlucCD9 and CD81KO-TlucCD9 cells. However, this enhancement was not observed in PanKO-TlucCD9 or PanKO-TlucCD63 cells. We initially hypothesized that silencing CD9 would boost engineered EV production in both CD9KO- and PanKO-cells upon reintroduction of TlucCD9. Instead, we observed that transduction with TlucCD9 led to expression or overexpression of other tetraspanins, indicating possible competition with exogenous TlucCD9. This competitive dynamic may limit engineered EV production. Additionally, the PanKO may impair key cellular processes, further affecting EV biogenesis. These findings suggest a need for further investigation into compensatory mechanisms among tetraspanins.

In contrast, reintroducing Tluc-CD81 revealed a different pattern. PanKO cells produced more engineered Tluc-EVs than CD81KO cells, suggesting that endogenous competition for Tluc-CD81 may be lower in PanKO cells. Moreover, Tluc-CD81 might fulfill essential physiological roles in PanKO cells, potentially enhancing EV release for intercellular signaling. When comparing KO lines, CD81KO cells released more engineered Tluc-EVs than CD9KO cells, though not significantly more than CD63KO cells. As shown in Figure 4E and 4F, introduction of Tluc-CD81 into CD9KO- and CD63KO-cells resulted in higher engineered EV production than in WT cells. This suggests that these KO backgrounds may face reduced competition during EV biogenesis, facilitating better incorporation of Tluc-CD81. Endogenous CD81 in CD9KO and CD63KO cells may still compete with exogenous CD81, but the reduced overall competition likely contributes to the enhanced EV yield.

To further explore this dynamic, we generated stably expressing CD63-mNG and CD81-mNG cell lines in both KO and WT backgrounds. Fluorescence data showed that all KO cell lines exhibited significantly higher EV production than WT controls. These results corroborate the luminescent assay data, providing consistent evidence that tetraspanin KO enhances engineered EV release when the corresponding tetraspanin is reintroduced. This cross-validation strengthens the conclusion that tetraspanins play key, context-dependent roles in EV biogenesis and represent promising targets for EV bioengineering strategies.

Lastly, to gain a deeper understanding, we performed proteomic analysis of CD63 in both CD63KO-Tluc-CD63 and WT-Tluc-CD63 EVs. The data provided strong evidence that CD63 could be fully restored and even elevated when endogenous CD63 was knocked out. This supports our hypothesis that there is competition between exogenous and endogenous tetraspanins. These findings are also consistent with our previous observations, where TSPAN2 and TSPAN3 were identified as the most effective for cargo loading compared to other tetraspanins when introduced into HEK-WT cells. This is likely because neither TSPAN2 nor TSPAN3 are highly expressed in EVs derived from wild-type HEK-293T cells[35].

## 4. Conclusion

In conclusion, our findings emphasize that while individual tetraspanins are not essential for EV biogenesis, their combined absence significantly reduces EV output. Reintroducing exogenous tetraspanins into PanKO-, CD9KO, CD63KO-, and CD81KO-cells enhances engineered EV production, particularly when using fusion constructs for protein loading. These results underline the critical roles of tetraspanins in natural EV biology and the optimization of EV-based therapeutic platforms.

## 5. Experimental Section

### Generation of monoclonal CD9/63/81 knockout cells

On Day 1, HEK293T cells were seeded at a density of 1 × 10⁴ cells per well in sterile, tissue culture-treated, flat-bottom 96-well plates containing complete Dulbecco’s Modified Eagle Medium (DMEM) (high glucose) supplemented with 10% fetal bovine serum (FBS; Gibco, USA) and 1% antibiotic-antimycotic solution (Invitrogen, USA). On Day 2, A total of 100 ng of Cas9 protein was used per well, corrisponding to approximately 5.5 nM Cas9 RNP. The RNP was formed by mixing Cas9 and sgRNA in a 1:1.1 molar ratio and complexed for 10 min at RT. Cas9/sgRNA ribonucleoprotein complexes were delivered to the cells using Lipofectamine™ RNAiMAX (Invitrogen) according to the protocol described in reference[36]. On Day 5, cells were detached using trypsin, then stained with fluorophore-conjugated antibodies targeting CD9, CD63, and CD81 (APC-conjugated; 2 µL per antibody), either individually or in combination. Staining was performed by incubating the cells with antibodies for 30 minutes at 4°C, followed by two washes with phosphate-buffered saline (PBS) to remove unbound antibodies. Subsequently, single cells exhibiting baseline tetraspanin expression were isolated using a SORP BD FACSAria™ Fusion cell sorter and deposited into receiving 96-well plates. Monoclonal cell populations were expanded and re-stained as described above, then analyzed via flow cytometry on a MACSQuant Analyzer 10 (Miltenyi Biotec) equipped with 405 nm, 488 nm, and 638 nm lasers. Clones demonstrating minimal tetraspanin expression were selected for further experimental use.

### Cell culture

HEK-293T wild-type (WT), single knockout (CD9KO, CD63KO, CD81KO), and triple tetraspanin knockout (PanKO) cell lines were maintained in complete DMEM as previously described. All cell lines were cultured in a humidified incubator at 37°C with 5% CO₂. Routine mycoplasma testing was conducted to ensure cell line integrity prior to EV isolation. For EV production, cells were seeded onto 15 cm culture dishes (Sarstedt, Germany) at approximately 60% confluency. After 24 hours, the growth medium was replaced with serum-free Opti-MEM (Gibco, USA), and cells were incubated for an additional 48 hours, by which time cultures reached approximately 90% confluency. Conditioned medium (CM) was subsequently harvested for EV purification.

### EV isolation

The CM was initially pre-cleared by centrifugation at 700 × g for 5 minutes to remove cells. Subsequently, to eliminate large cell debris and apoptotic bodies, the CM underwent a second centrifugation step at 2,000 × g for 10 minutes. The supernatant was then filtered through low protein-binding bottle-top filters (Corning) or 0.22 µm pore size cellulose acetate syringe filters (VWR, Sweden) to exclude residual large vesicles. EV concentration was performed using tangential flow filtration (TFF) on a KrosFlo Research 2i system (Spectrum Labs), operating at a flow rate of 100 mL/min and a transmembrane pressure of approximately 3 psi. A 300 kDa polyether sulfone hollow fiber filter (MidiKros, 370 cm² surface area; Spectrum Labs) was employed for EV retention. The sample was diafiltered with phosphate-buffered saline (0.1 M PBS) through 0.22 µm filters (Nalgene Rapid-Flow; Thermo Fisher Scientific, USA) using a volume equivalent to twice the original CM, resulting in a final concentrate volume of 20–35 mL.

### Nanoparticle tracking analysis (NTA)

Particle size distribution and concentration were measured using a NanoSight NS500 instrument (Malvern Panalytical) equipped with NTA 2.3 analytical software and a 488 nm laser. Prior to analysis, samples were diluted in 0.22 µm filtered PBS to achieve a particle concentration within the optimal detection range of 40 to 100 particles per frame. For each sample, up to five video recordings of 30 seconds each were captured in light scatter mode, with the camera level set between 13 and 15. Instrument settings, including screen gain (10) and detection threshold (7), were maintained consistently across all measurements.

### Measurement of luciferase Relative Luminescence Units (RLU) for engineered Tluc-EVs

Purified EVs (30 µL) isolated from conditioned media of each cell line expressing different Tluc constructs were lysed using 0.1% Triton X-100 (Sigma-Aldrich, USA). Lysates were transferred to white-walled 96-well plates, followed by the addition of 30 µL luciferin substrate (Firefly Luciferase Assay System; Promega) according to the manufacturer’s protocol. Luminescence was immediately measured using a GloMax® 96 Microplate Luminometer (Promega, USA). Luciferase activity was reported as relative light units (RLU) normalized to the number of EVs.

### Single-vesicle imaging flow cytometry for engineered mNG-EVs

Single-vesicle imaging flow cytometry was conducted using an Amnis Cellstream instrument (Amnis/Cytek, USA). Briefly, 2.5 × 10^8^ EVs, quantified by NTA, were stained with 8 nM fluorescent antibodies targeting the tetraspanins CD9-APC (clone: SN4 C3-3A2, cat. 130-128-037), CD63-APC (clone: H5C6, cat. 130-100-182, Miltenyi Biotec), and CD81-APC (clone: JS64, Beckman Coulter). The EVs were incubated overnight at room temperature in the dark[37]. Following this incubation, the concentration was diluted to 1 × 10^7^ EVs particles per ml in a final volume of 100 µl for data acquisition. Control samples included unstained EVs, and non-EV-containing samples incubated with the antibodies. PBS-HAT buffer[38] was used as the diluent for all steps. The results were analyzed using FlowJo software (v. 10.7.2; FlowJo LLC).

### Cell surface expression of tetraspanins

All cell lines were cultured for three days in complete DMEM as described above. 2×10^5^ cells were added into V-bottom 96-well plates and stained with either APC (allophycocyanin)-conjugated anti-human (clone: SN4 C3-3A2, cat. 130-128-037), CD63-APC (clone: H5C6, cat. 130-100-182, Miltenyi Biotec), and CD81-APC (clone: JS64, Beckman Coulter) at a concentration on 1µg/ml at 4 °C for 30 min. Then, the plate was spun at 900×g for 5 minutes to remove free antibodies. The cells were resuspended in 100 µl of PBS and stained with 0.1µg/ml DAPI (4′, 6-diamidino-2-phenylindole). Finally, the plate was run by a MACSQuant 10 Flow Cytometer (Miltenyi Biotec, USA), and the data was analyzed by using FlowJo software (v. 10.7.2; FlowJo LLC).

### Surface expression of tetraspanins on the EVs

50 µl of purified EVs at a concentration of 1 × 10^10^/ml were stained with either anti-human CD9 (Miltenyi Biotech, clone SN4), anti-human CD63 (Miltenyi Biotec, clone H5C6) and anti-human CD81 antibodies (Beckman Coulter, clone JS64) or REA control APC conjugated antibodies at a concentration of 8 nM overnight at room temperature in the dark. The EVs were diluted to a concentration of 5 × 10^6^/mL to detect the surface expression of tetraspanins using an Amnis CellStream Flow Cytometer (Luminex, Cytek, USA), and the data was analyzed by using FlowJo software (v. 10.7.2; FlowJo LLC).

### Transmission Electron Microscopy (TEM)

Three microliters of the sample were applied to glow-discharged, carbon-coated, formvar-stabilized 400 mesh copper grids (easiGlow™, Ted Pella) and incubated for approximately 30 seconds. The excess sample was then blotted off, followed by a MilliQ water wash. Grids were negatively stained with 1% ammonium molybdate and imaged using an HT7800-Xarosa transmission electron microscope (Hitachi High-Technologies) operated at 80 kV, equipped with a 4 MP Veleta CCD camera (Olympus Soft Imaging Solutions GmbH).

### Lentiviral supernatant production

HEK-293T cells were seeded at a density of 22 × 10⁶ cells per T175 tissue culture flask (Sarstedt, Germany) in complete DMEM and maintained in a humidified incubator at 37°C with 5% CO₂ for 24 hours. On the following day, the medium was carefully aspirated and replaced with 15 mL of fresh complete DMEM. For transfection, 22 µg of each plasmid construct (Tluc-CD9-Cerulean, Tluc-CD63-Cerulean, Tluc-CD81-Cerulean, CD63-mNG, and CD81-mNG) was combined with 3.5 µg of envelope plasmid pcoPE, 22 µg of helper plasmid pCD/NL-BH, and 27 µg of polyethyleneimine (PEI) in 3 mL of Opti-MEM (serum- and antibiotic-free). The transfection mixture was then added to the cells, which were incubated for 22 hours. On Day 3, the medium was replaced with 20 mL of complete DMEM supplemented with 10 mM sodium butyrate (Sigma-Aldrich, USA) to enhance expression via the human CMV immediate early promoter. After 6–7 hours, the medium was exchanged for 20 mL of fresh complete DMEM, and the cells were incubated for an additional 18–22 hours. On Day 4, the conditioned medium was collected, filtered through a 0.45 µm syringe filter (VWR, Sweden), and subjected to ultracentrifugation at 25,000 × g for 90 minutes at 4°C to pellet viral particles. The viral pellets were resuspended in 1 mL of Iscove’s Modified Dulbecco’s Medium (IMDM) supplemented with 1% Antibiotic-Antimycotic (Invitrogen, USA) and 20% fetal bovine serum (FBS), and aliquots were stored at −80°C until use.

### Generation of stable cell lines

The viral particles were titrated before creating stable cell lines to determine the necessary dosages. 1×10^5^ HEK293T WT, PanKO, CD9KO, CD63KO, and CD81KO cells were seeded in complete DMEM on 12-well plates (Sarstedt, Germany) and incubated 24 hours prior to transduction. The cells were transduced at MOI of 5 for 20 to 24 hours. The medium from the cells was aspirated gently and replaced with a new complete DMEM. Subsequently, the cells underwent at least five rounds of selection using 1000 µg/mL Zeocin or 4 µg/mL Puromycin (both Thermo Fisher Scientific, USA). Cell growth was monitored throughout the procedure, with regular passaging using a selection antibiotic before the cell sorting step.

### Fluorescence-activated cell sorting (FACS)

Once selected, the cells were sorted using a MA900 Multi-Application Cell Sorter (Sony Biotechnology, USA). Cerulean was detected using 433 nm excitation and 475 nm emission, while mNG was detected at 488 nm. After sorting, the cell with the highest fluorescence signal of interest was selected. At the same time, 100,000 cells from each line were sorted into 12-well plates with sorting buffer. The cells were then cultured in complete DMEM in a 37°C, 5% CO₂ humidified incubator.

### Plasmid constructs

Human CD9, human CD63, and human CD81 sequences were optimized using the UniProt database and the luciferase proteins as published before[28]. Gene segments (Integrated DNA Technologies) encoding the CD9, CD63, CD81, and luciferase proteins were cloned using NotI and EcoRI into the pLEX vector backbone containing a CAG promoter. Using NotI and SacI, we generated distinct plasmids to produce the luciferase protein Tluc linked to the N terminus of CD9, CD63, and CD81 in the pLEX vector. Tluc was fused to CD9, CD63, and CD81 to create the C terminus using BsiWI and EcoRI in the pLEX vector. The DNA codon optimization for Cerulean was confirmed by the Uniprot database and cloned into lentiviral vector p2CL9IPwo5[39]as a backbone downstream of the SFFV promoter using SbfI and BsrGI and upstream of an internal ribosomal entry site-zeocin resistance cDNA cassette for the generation of Cerulean constructs. Then, using NotI and EcoRI, all N-terminal Tluc constructs, Tluc-CD9, Tluc-CD63, and Tluc-CD81, were subcloned into a Cerulean construct. Sequencing was used to validate all designs. For CD63-mNG and CD81-mNG cloned into p2CL9IPw5 vector using NotI and EcoRI.

### Cell proliferation assay

A total of 15 × 10³ cells from Pan-KO, CD9-KO, CD63-KO, CD81-KO, and WT HEK-293T cell lines were seeded into individual wells of a 96-well plate containing complete DMEM. Cells were incubated for 24 hours at 37 °C in a humidified atmosphere with 5% CO₂. Following the incubation period, 10 μL of WST-1 reagent (Merck KGaA, Darmstadt, Germany) was added to each well and incubated for either 30 minutes or 4 hours at 37 °C in 5% CO₂, in accordance with the manufacturer’s instructions. Absorbance was subsequently measured at 440 nm using a microplate spectrophotometer (SpectraMax® i3x; Molecular Devices LLC, USA).

### Transfection

All cell types were seeded into 96-well plates at a density of 2 × 10⁴ cells per well, 24 hours prior to transfection. For each transfection, 100 ng of plasmid DNA encoding eGFP-tagged CD9, CD63, or CD81 was diluted in 10 μL of Opti-MEM and incubated for 5 minutes at room temperature. In parallel, 2 μg of Lipofectamine 2000 (Sigma) was diluted in 10 μL of Opti-MEM (maintaining a 1:2 DNA to reagent ratio) and also incubated for 5 minutes at room temperature. The diluted DNA and Lipofectamine solutions were then combined, gently mixed, and incubated for an additional 25 minutes at room temperature to facilitate complex formation. The resulting transfection complexes were added dropwise to the corresponding wells. After 24 hours, cells were trypsinized, resuspended in complete DMEM, and analyzed using a MACSQuant 10 flow cytometer (Miltenyi Biotec, USA). Data acquisition was followed by analysis using FlowJo software (version 10.7.2; FlowJo LLC).

### Bind-elute and size exclusion (BE-SEC) chromatography

The pre-concentrated CM was subjected to size-exclusion chromatography using BE-SEC columns (HiScreen Capto Core 700, GE Healthcare Life Sciences) integrated with the ÄKTA Pure 25 chromatography system (GE Healthcare Life Sciences)[40]. Column equilibration, sample loading, and column cleaning procedures (CIP) were performed in accordance with the manufacturer’s guidelines, with appropriate flow rates applied at each step. EV fractions were collected based on UV absorbance profiles at 280 nm. The collected fractions were subsequently washed with 30 mL of PBS filtered through a 0.22 µm membrane, followed by concentration using Amicon Ultra-15 centrifugal filters with a 10 kDa molecular weight cut-off (Millipore) to a final volume of 300 μL. The concentrated EV samples were then stored at −80 °C for subsequent protein quantification.

### Protein quantification of the cells

The cells (1×10^5^) were pelleted at 500×g for 5 minutes at 4°C and resuspended in cold 0.22µm filtered PBS and re-pelleted. The pellets were then lysed in 100µl of 0.1% Triton X-100 (Sigma-Aldrich, USA) followed by protein quantification with the DC Protein Assay kit (Bio-Rad).

### Proteomics Analysis

Prior to proteomic analysis, protein concentrations of EV samples were determined using the Micro BCA™ Protein Assay Kit (Thermo Fisher Scientific, USA) according to the manufacturer’s instructions. EVs derived from WT, KO, or stably transduced cell lines were subjected to liquid chromatography-tandem mass spectrometry (LC-MS/MS) analysis. Briefly, samples were analyzed using a Thermo Fisher Scientific Q Exactive Plus system employing a 1-hour chromatographic gradient. Data processing and statistical analysis were conducted using R (version 4.1.2) and RStudio (version 2022.07.1+554), utilizing the DEP package (version 1.16.0). Proteins uniquely of bovine origin, together with other contaminants (e.g., Tluc, Cerulean, trypsin, keratins) were excluded from the dataset prior to analysis. Following filtering, 2,355 proteins were retained across all samples for downstream analysis. Differentially enriched proteins were further analyzed by Gene Ontology (GO) enrichment and classified according to Protein Ontology categories using the PANTHER classification system (version 17.0). The mass spectrometry proteomics dataset has been deposited in the ProteomeXchange Consortium via the PRIDE repository.

### STRING Analysis

The STRING database (https://string-db.org) was used to make protein–protein interaction networks for CD9, CD63, and CD81. The analysis was conducted using default parameters, employing a high-confidence interaction score (≥0.7). Partners in the interaction were only those that had been experimentally tested and had been added to a database. The resulting network depicted anticipated interactions among CD9, CD63, and CD81, along with other members of the tetraspanin family.

### Statistical analysis

Data are expressed as mean ± standard deviation (SD). Statistical comparisons were conducted using one-way analysis of variance (ANOVA) followed by Tukey’s post hoc multiple comparisons test. A p-value < 0.05 was considered indicative of statistical significance. All analyses were performed using GraphPad Prism version 10.

## Acknowledgements

Oscar P. B. Wiklander discloses support for the research described in this study from the The Swedish Research Council (VR, 2022-02449), The Swedish Cancer Society (Cancerfonden, project 23 2935 Pj), Radiumhemmet (project #241392), the Center for Innovative Medicine (CIMED) junior investigator grants (FoUI-976434), and Karolinska Institutet (2-116/2023). Samir E.L. Andaloussi discloses support for the research described in this study from Horizon 2020 (EXPERT, 4-2298/2019), the Swedish Foundation of Strategic Research (FormulaEx, SM19-0007), European Research Council Consolidator Grant (DELIVER, 101001374), the Swedish Cancer Society (4-511/2022) and the Swedish Research Council (4–258/202). Joel Z. Nordin discloses support for the research described in this study from the Swedish Research Council (VR) (2021-02407) and the CIMED junior investigator grant. Helena Sork discloses support from the Estonian Research Council grants (PSG1043 and PRG1882) and NEURON JTC2023 partnership, co-funded by the European Union and Estonian Research Council through the Mobilitas 3.0 (MOB3ERA6). We would like to acknowledge the MedH Flow Cytometry core facility (Karolinska Institutet), supported by KI/SLL, for providing cell sorting services.

## Author contributions

Doste R. Mamand: Conceptualization, data curation, formal analysis, investigation, methodology, resources, supervision, visualization, writing—original draft, writing—review, and editing. Helena Sork: Data curation, formal analysis, investigation, methodology, validation, visualization. Rim Jawad Wiklander: Formal analysis, investigation, methodology. Safa Bazaz: Data curation, formal analysis, investigation, methodology. Oskar Gustafsson: Data curation, formal analysis, investigation, methodology, validation. Xiuming Liang: Investigation, methodology. Dhanu Gupta: Investigation, methodology, writing—review and editing. Vicky W.Q. Hou: Investigation, writing—review and editing. André Görgens: Formal analysis, investigation, methodology, writing—review and editing. Joel Z. Nordin: Formal analysis, investigation, methodology, writing—review and editing. Samir El-Andaloussi: Conceptualization, data curation, formal analysis, funding acquisition, investigation, methodology, project administration, resources, supervision, writing—review, and editing. Oscar P. B. Wiklander: Conceptualization, data curation, formal analysis, investigation, methodology, supervision, visualization, writing—review, and editing.

## Data Availability Statement

The data that support the findings of this study are available from the corresponding author upon reasonable request.

