## supplementary for "Evaluation of Tetraspanins in Extracellular Vesicle Bioengineering": Revised Supplementary Data and Figures. BioRixv.pdf

*Correspondence

[#]These authors contributed equally

Supplementary Figure 1

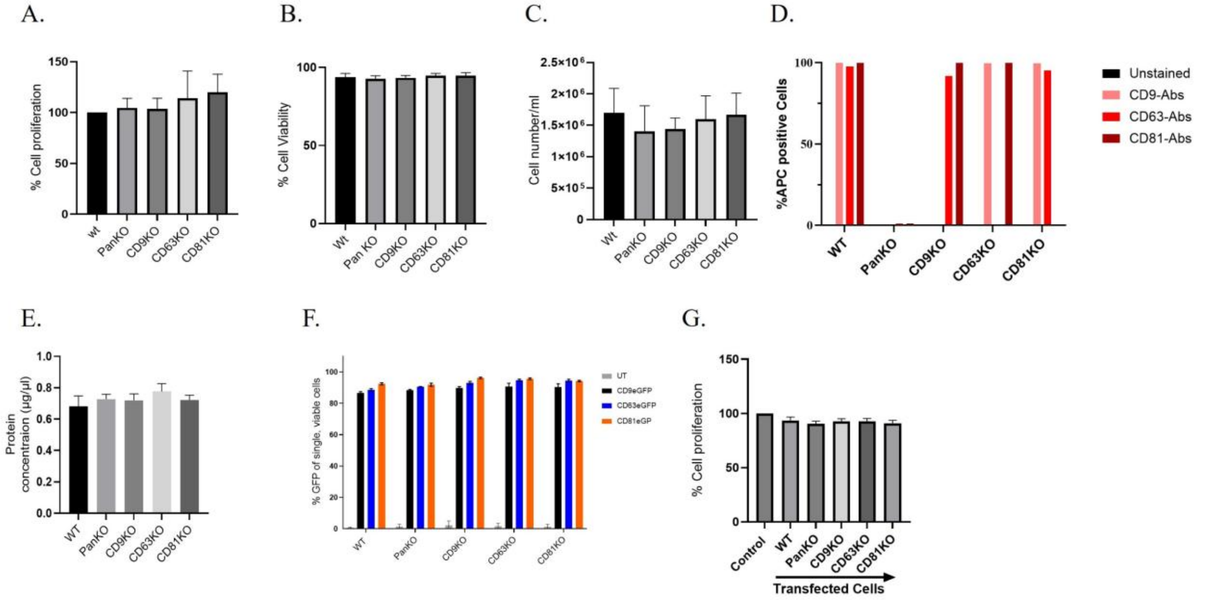


**Supplementary Figure 1**: Characterization of the cell lines after single-cell knockout tetraspanins. A: Cell proliferation assay by performing WST-1 at absorbance of 450nm to 650nm. B. Cell viability assay using trypan blue. C: Cell counting for each cell line (cells/ml) when reaching confluency using cell Countess. D. Quantification of FC data for WT and KO cells. E. DC Protein Assay measurement at $1\times10^5$ cells for each cell line. F. Transfection efficiency of the cells by Lipofectamine 2000 with CD9eGFP, CD63eGFP, and CD81eGFP plasmids. G. Cell proliferation assay for the transfected cell by performing WST-1 at absorbance of 450nm to 650nm. The data are presented as means (±SD, n = 3). One way ANOVA was used for statistical analysis.

### Supplementary Figure 2

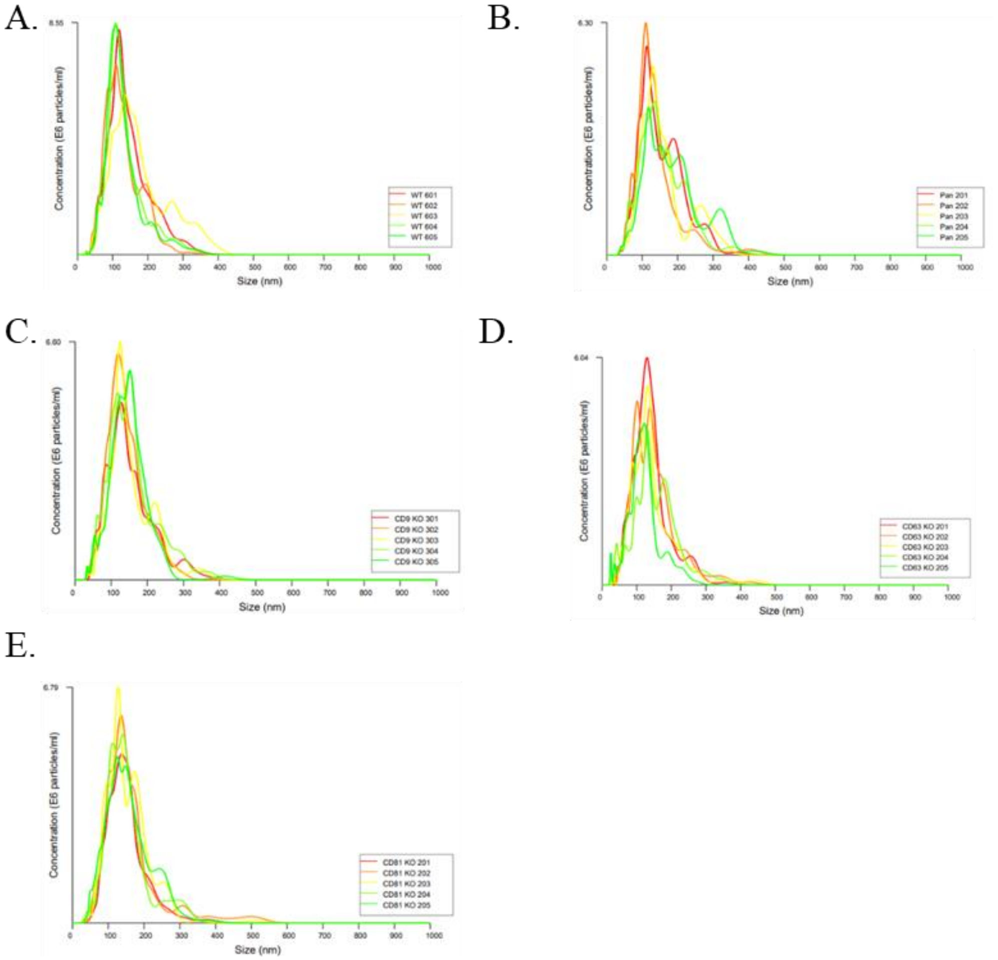


**Supplementary Figure 2**. Detection of EV size distribution using NTA. A: Size of WT-EVs in nm. B: Size of PanKO-EVs in nm. C: Size of CD9KO-EVs in nm. D: Size of CD63KO-EVs in nm. E: Size of CD81KO-EVs in nm. NanoSight NS500 equipped with NTA 3.2 analytical software (Malvern Panalytic, UK) was used.

Supplementary Figure 3

A. WT-EVs

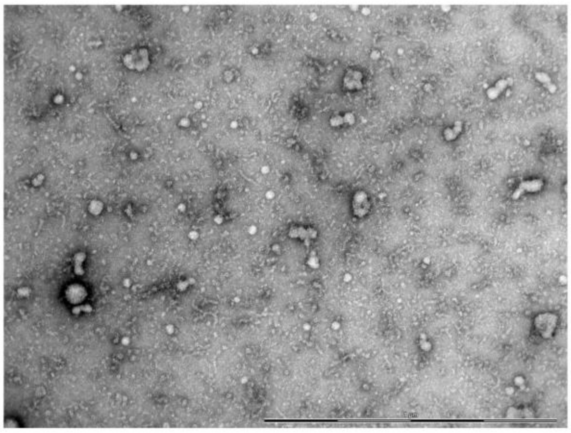

B. PanKO-EVs

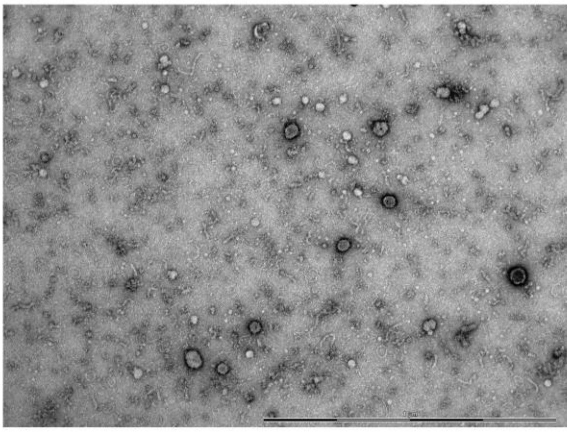

C. CD9-EVs

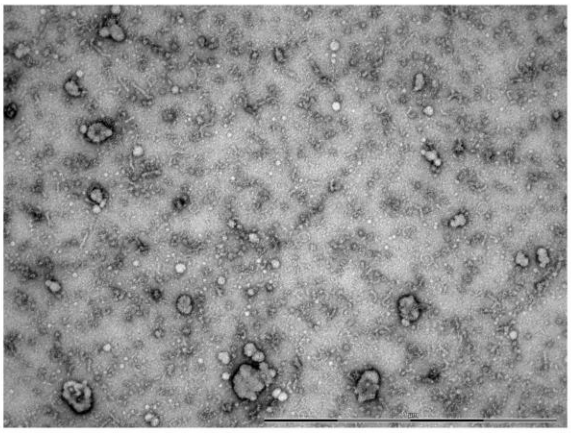

D. CD63KO-EVs

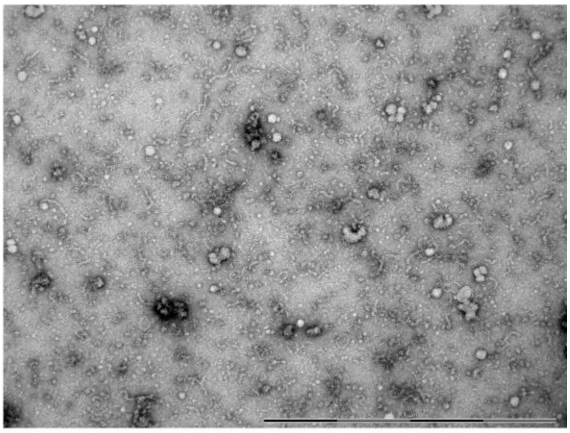

E. CD81KO-EVs

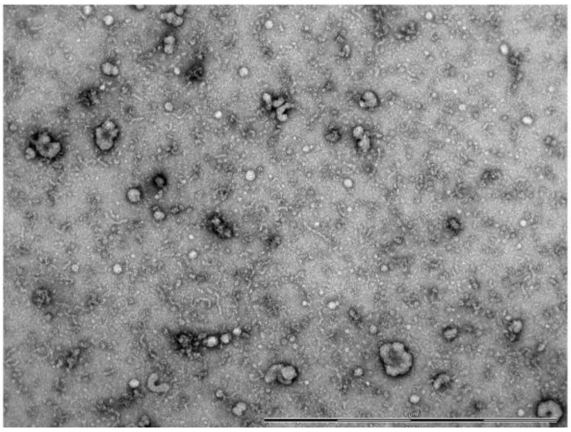

Supplementary Figure 3. Detection of EVs by TEM. Representative images of. A: WT-EVs. B: PanKO-EVs. C: CD9KO-EVs. D: CD63KO-EVs. E: CD81KO-EVs. Scale bar: 1 μm.

**Table 1**: Generating cells lines stably expressing Cerulean Tluc-CD9 or Tluc-CD63 or Tluc-CD81.

| No. | Not Engineered cells | Stably expressing Tluc-CD9 | Stably expressing Tluc-CD63 | Stably expressing Tluc-CD81 |
|---|---|---|---|---|
| 1 | HEK293T-WT | HEK293T-WT | HEK293T-WT | HEK293T-WT |
| 2 | HEK293T-PanKO | HEK293T-PanKO | HEK293T-PanKO | HEK293T-PanKO |
| 3 | HEK293T-CD9KO | HEK293T-CD9KO | HEK293T-CD9KO | HEK293T-CD9KO |
| 4 | HEK293T-CD63KO | HEK293T-CD63KO | HEK293T-CD63KO | HEK293T-CD63KO |
| 5 | HEK293T-CD81KO | HEK293T-CD81KO | HEK293T-CD81KO | HEK293T-CD81KO |

Supplementary Figure 4

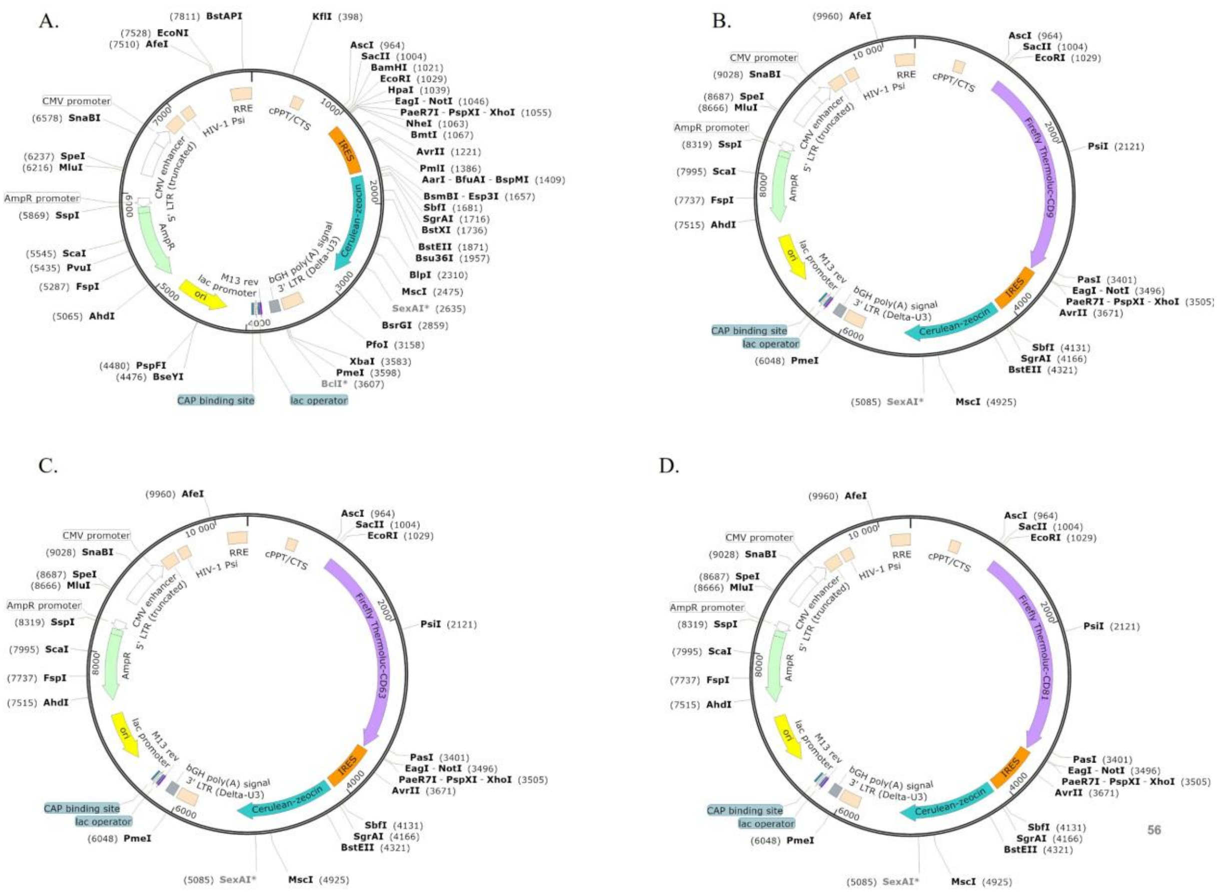


**Supplementary Figure 4**: Plasmid map for Cerulean expression constructs. A: Schematic illustration of Cerulean expression lentiviral constructs as internal control. B: Schematic illustration of lentiviral construct expressing Tluc-CD9 not tagged to Cerulean. C: Schematic illustration of lentiviral construct expressing Tluc-CD63 not tagged to Cerulean D: Schematic illustration of lentiviral construct expressing Tluc-CD81 not tagged to Cerulean.

Supplementary Figure 5

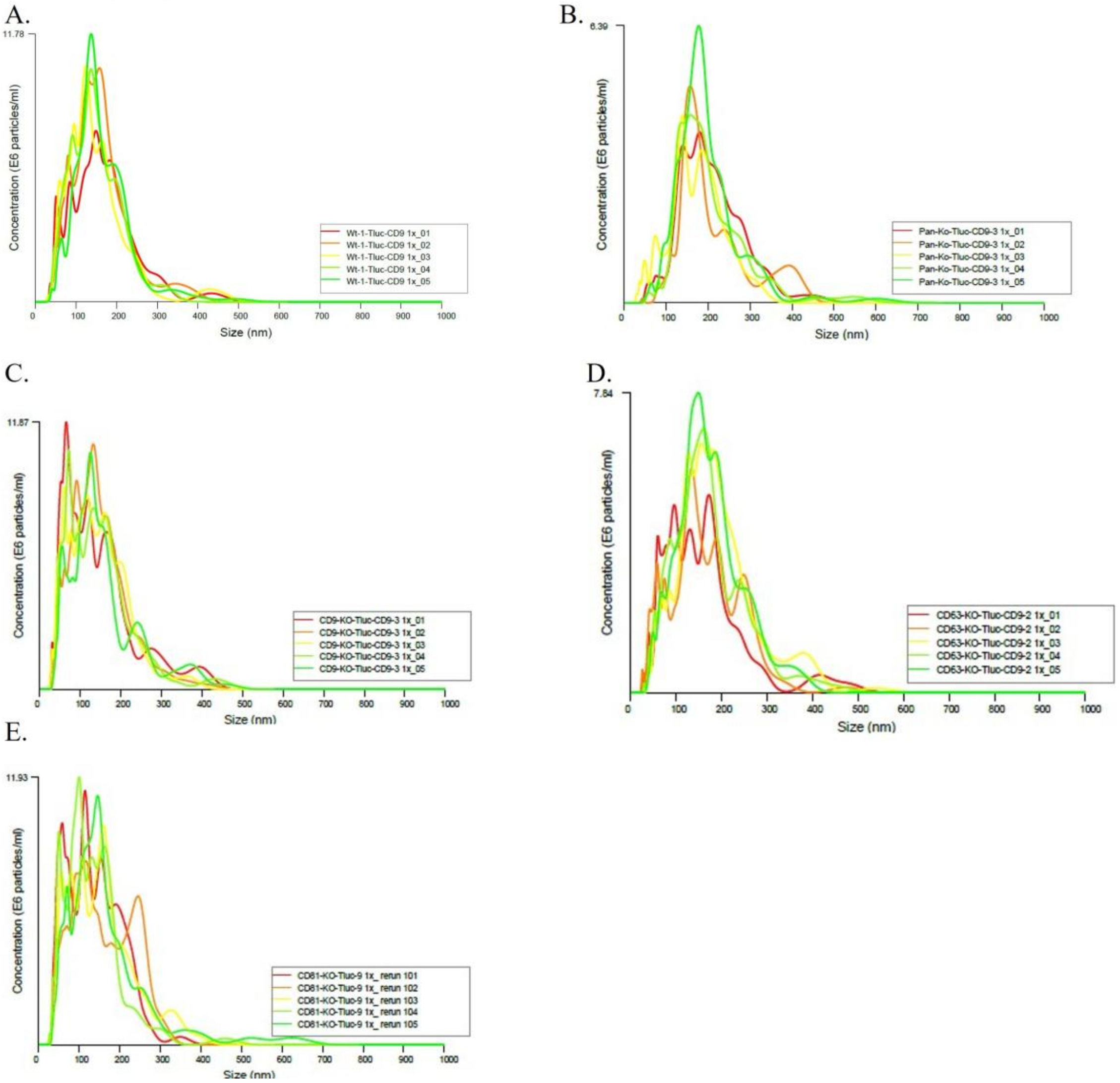


**Supplementary Figure 5**. Detection of EV size distribution using NTA. A: Size of WT- Tluc-CD9EVs nm. B: Size of PanKO- Tluc-CD9EVs in nm. C: Size of CD9KO- Tluc-CD9EVs in nm. D: Size of CD63KO- Tluc-CD9EVs in nm. E: Size of CD81KO- Tluc-CD9EVs in nm. NanoSight NS500 equipped with NTA 3.2 analytical software (Malvern Panalytic, UK) was used.

Supplementary Figure 6

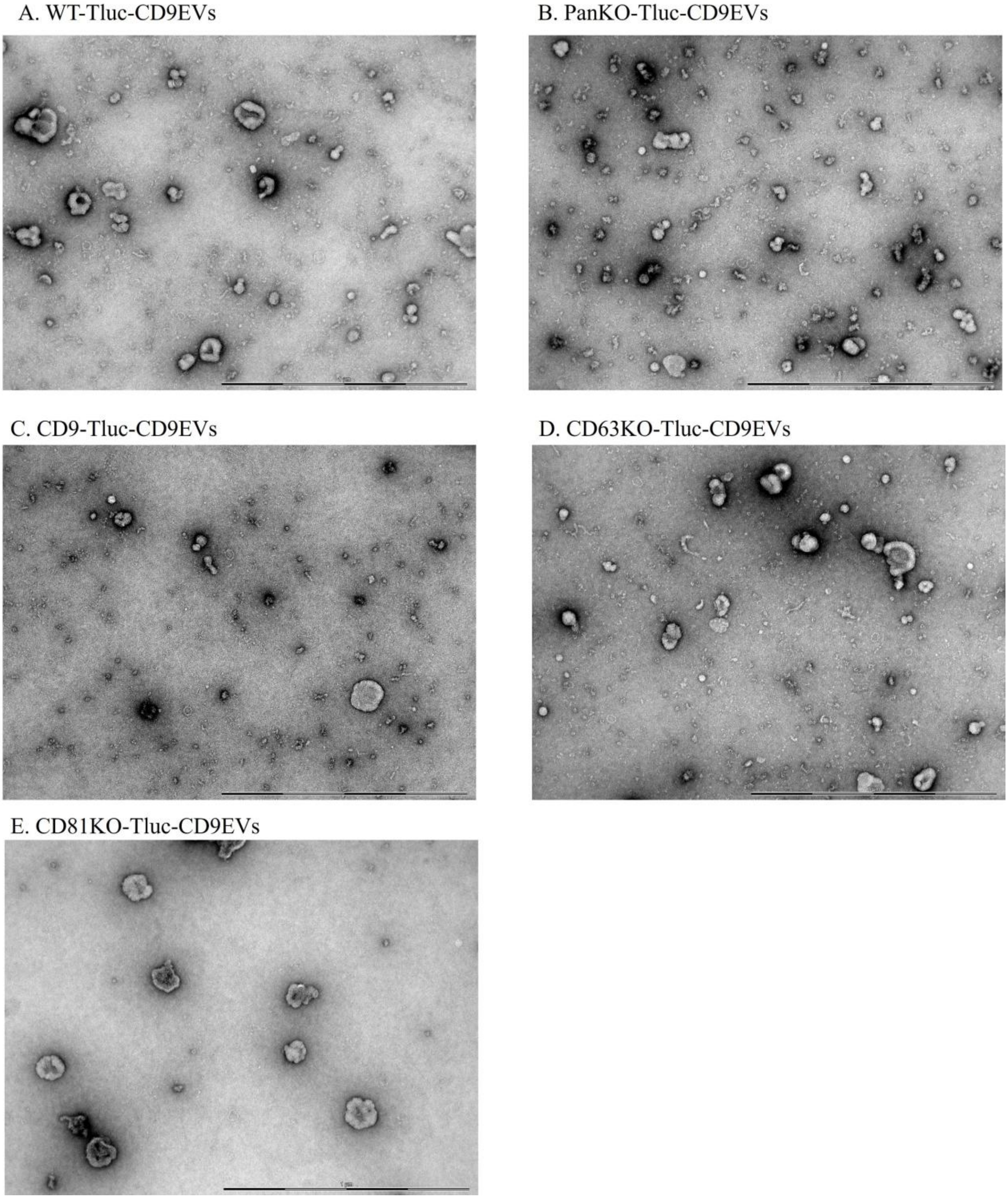


**Supplementary Figure 6**. Detection of EVs by TEM. Representative images of. A: WT-Tluc-CD9EVs. B: PanKO- Tluc-CD9EVs. C: CD9KO- Tluc-CD9EVs. D: CD63KO- Tluc-CD9EVs. E: CD81KO- Tluc-CD9EVs. Scale bar: 1 μm.

Supplementary Figure 7

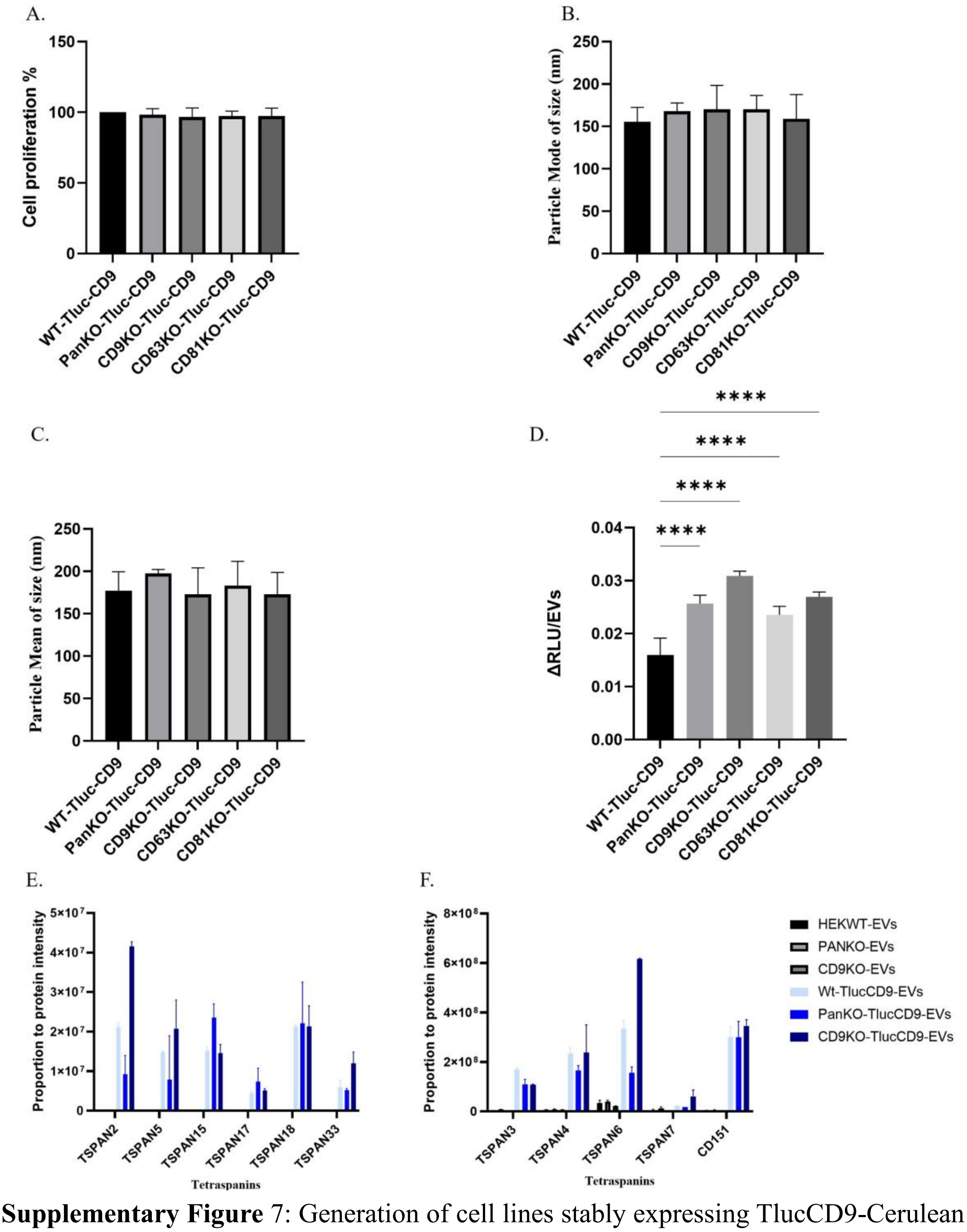


**Supplementary Figure** 7: Generation of cell lines stably expressing TlucCD9-Cerulean using lentivirus transduction of WT, PanKO, CD9KO, CD63KO, and CD81KO cells. A. Cell proliferation assay by performing WST-1 at absorbance of 450nm to 650nm. B. Particle mode of size in nm. C. Particle mean of size in nm. D . Relative luminescence Unit (RLU) of engineered Tluc-CD9-EVs produced by KO cells and WT cells (normalized over free RLU). E and F. Expression and overexpression of tetraspanins from proteomics analysis using MaxQuant data-driven mass spectrometry for EVs from KO and WT cells (TOTAL MS AREA/tetraspanin MS Area). The data are presented as means (±SD, n = 3-6). One-way ANOVA was used to show significance and was illustrated as follows: **** $p < 0.0001$.

Supplementary Figure 8

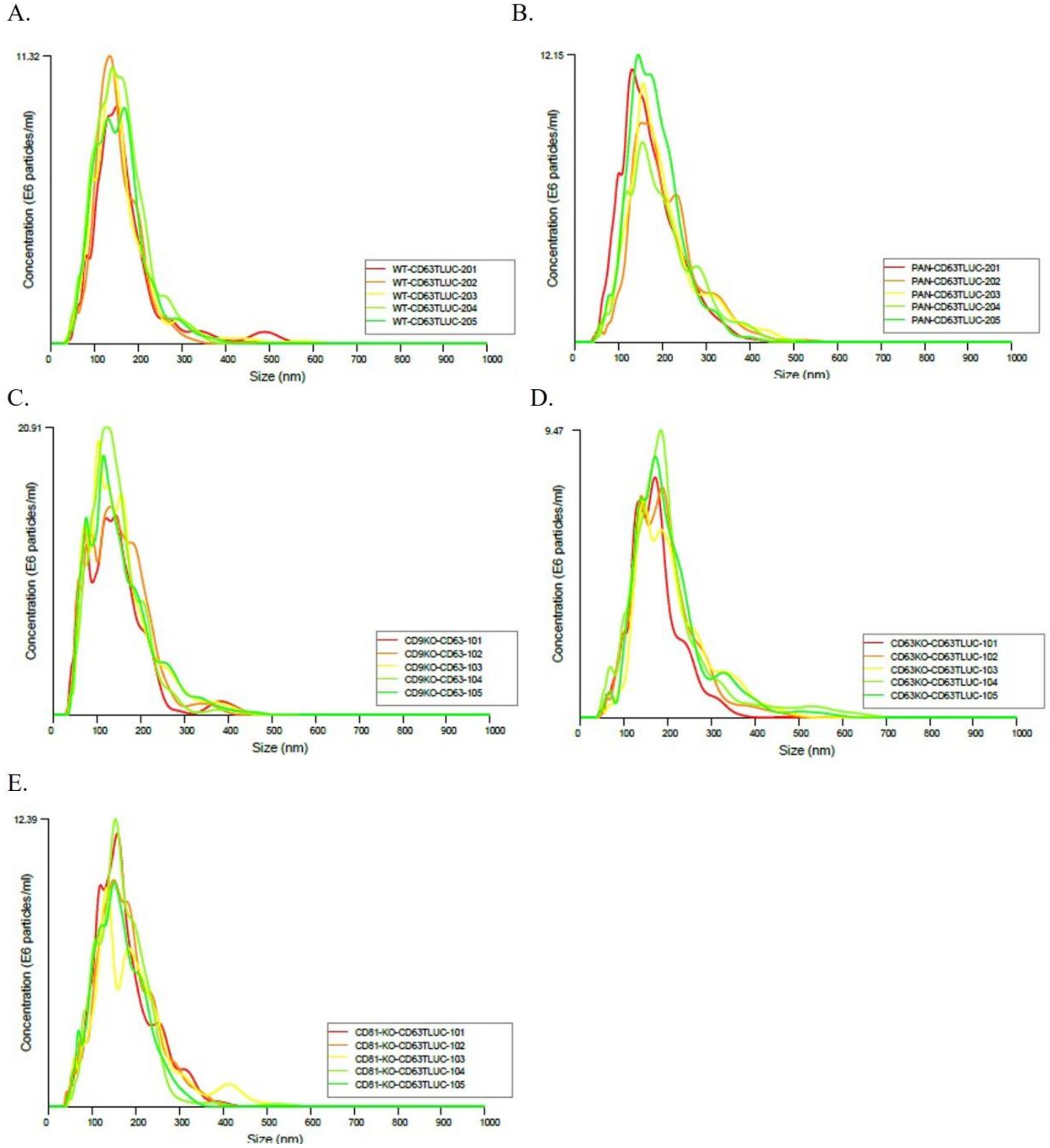


**Supplementary Figure 8**. Detection of EV size distribution using NTA. A: Size of WT- Tluc-CD63EVs nm. B: Size of PanKO-Tluc-CD63EVs in nm. C: Size of CD9KO- Tluc-CD63EVs in nm. D: Size of CD63KO-Tluc-CD63EVs in nm. E: Size of CD81KO-Tluc-CD63EVs in nm. NanoSight NS500 equipped with NTA 3.2 analytical software (Malvern Panalytic, UK) was used.

Supplementary Figure 9

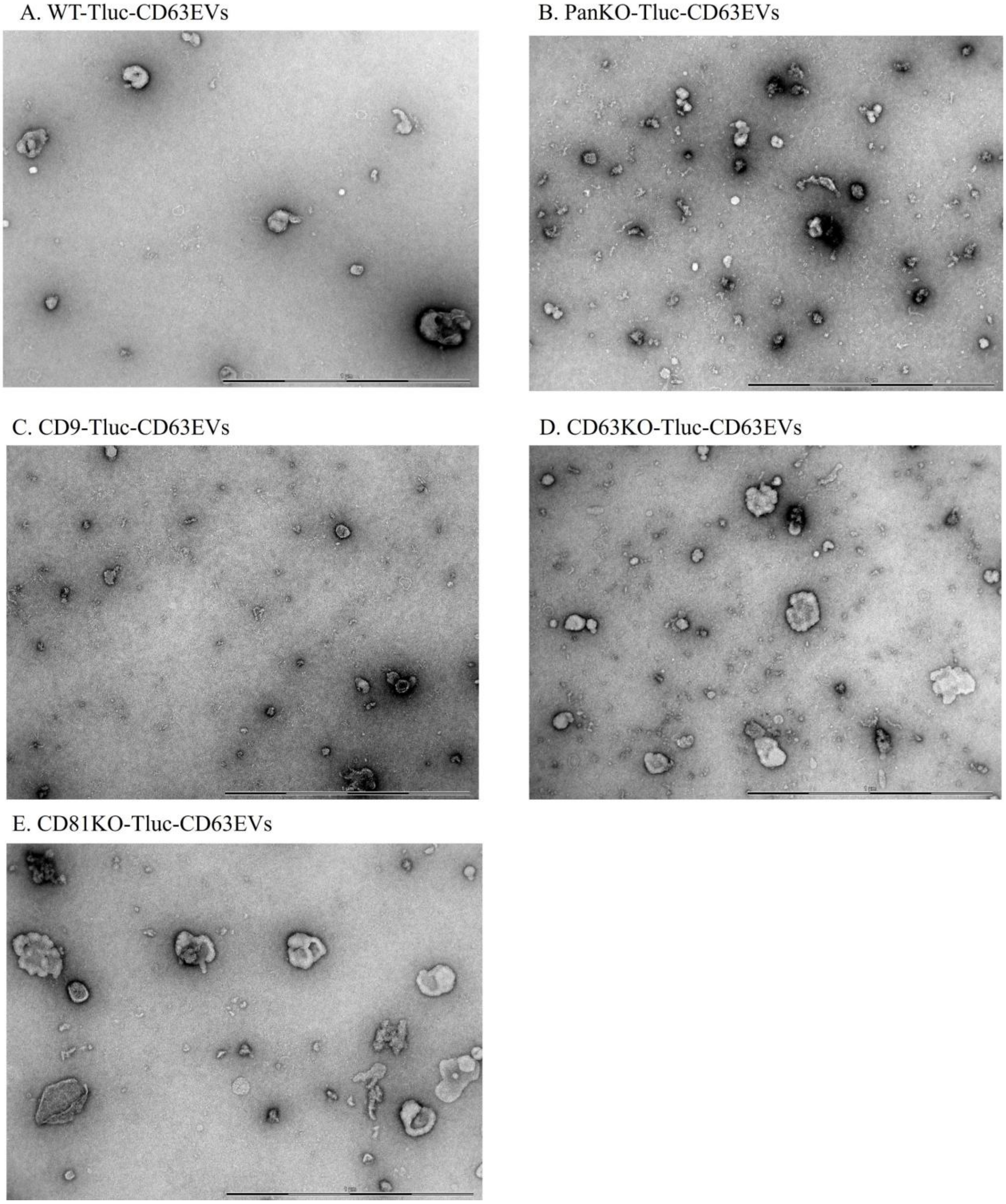


**Supplementary Figure 9**. Detection of EVs by TEM. Representative images of. A: WT-Tluc-CD63EVs. B: PanKO- Tluc-CD63EVs. C: CD9KO- Tluc-CD63EVs. D: CD63KO-Tluc-CD63EVs. E: CD81KO- Tluc-CD63EVs. Scale bar: 1 µm.

Supplementary Figure 10

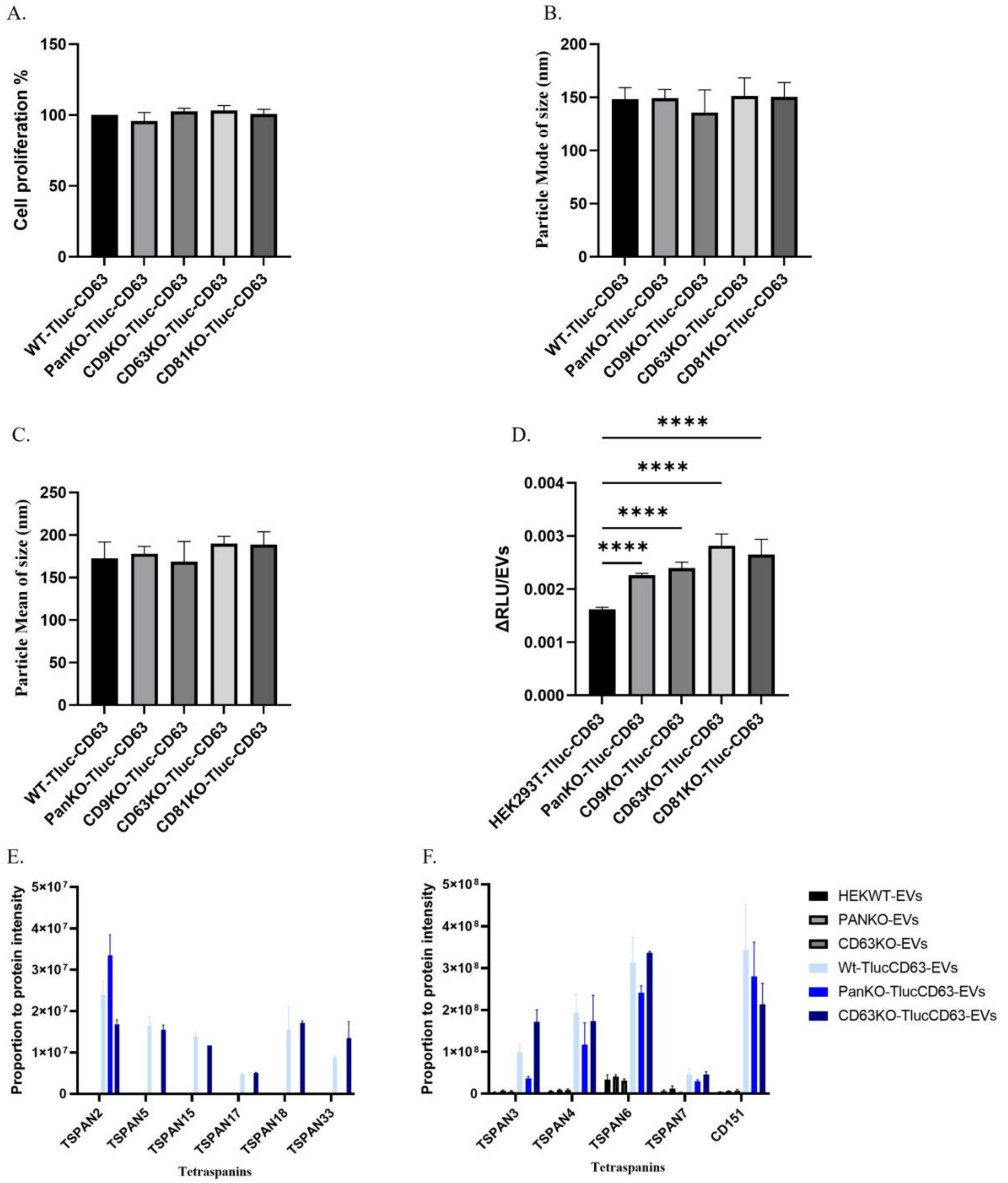


**Supplementary Figure 10**: Generation of stably expressing TlucCD63-Cerulean by lentiviruses in WT, PanKO, CD9KO, CD63KO, and CD81KO cells. A. Cell proliferation assay by performing WST-1 at absorbance of 450nm to 650nm. B. Particle mode of size in nm. C. Particle mean of size in nm. D . Relative luminescence unit (RLU) of engineered Tluc-CD63-EVs produced by KO cells and WT cells (normalized over free RLU). E and F. Expression and overexpression of tetraspanin from proteomics using MaxQuant data-driven mass spectrometry for EVs from KO and WT cells (TOTAL MS AREA/tetraspanin MS Area). The data are presented as means (±SD, n = 3-6). One-way ANOVA was used to show significance and was illustrated as follows: **** $p < 0.0001$.

Supplementary Figure 11

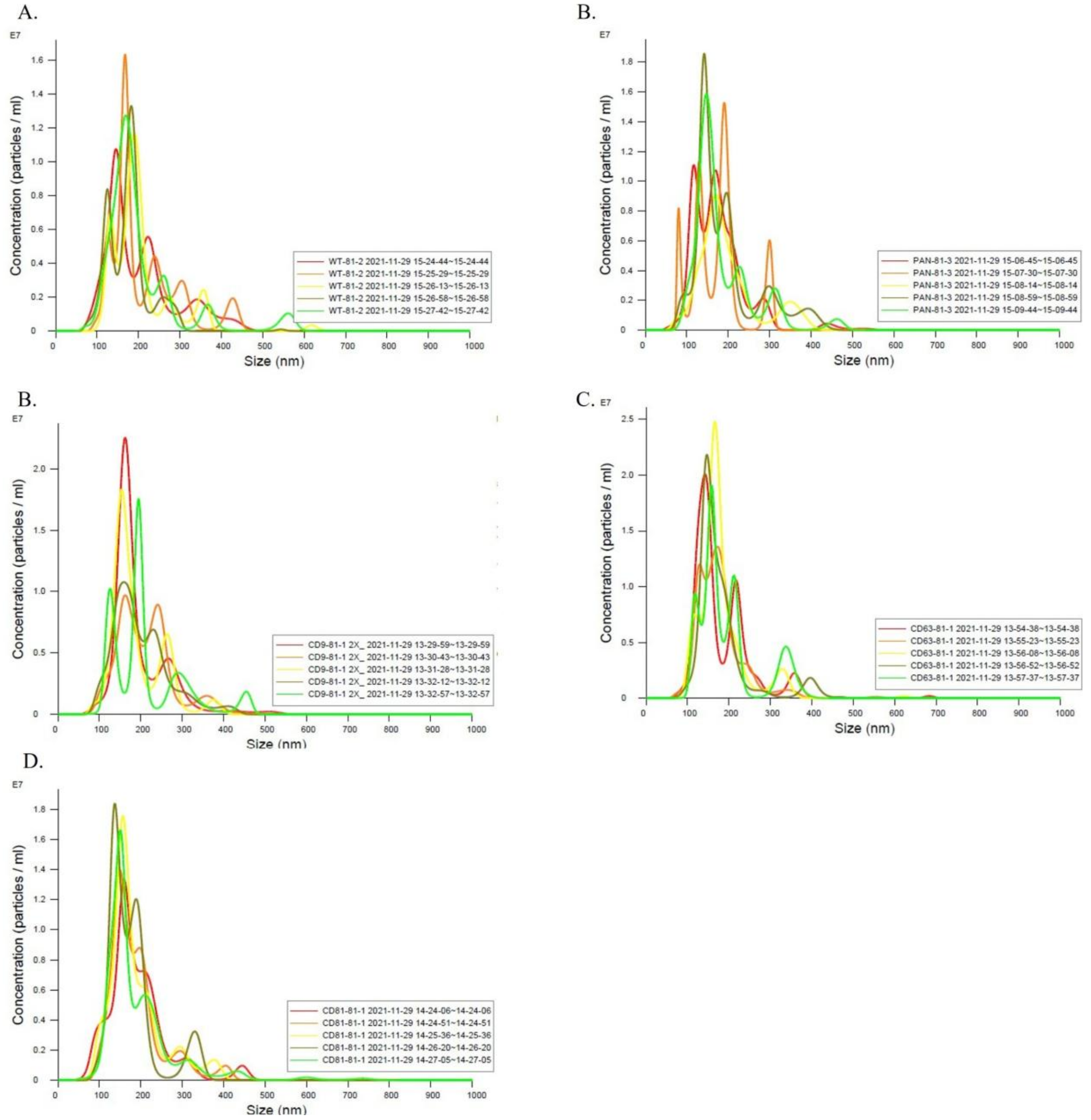


**Supplementary Figure 11**. Detection of EV size distribution using NTA. A: Size of WT- Tluc-CD81EVs nm. B: Size of PanKO-Tluc-CD81EVs in nm. C: Size of CD9KO- Tluc-CD81EVs in nm. D: Size of CD63KO- Tluc-CD81EVs in nm. E: Size of CD81KO-Tluc-CD81EVs in nm. NanoSight NS500 equipped with NTA 3.2 analytical software (Malvern Panalytic, UK) was used.

Supplementary Figure 12

A. WT-Tluc-CD81EVs

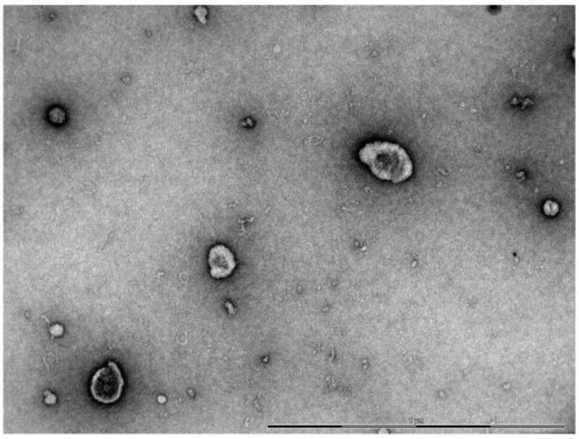

B. PanKO-Tluc-CD81EVs

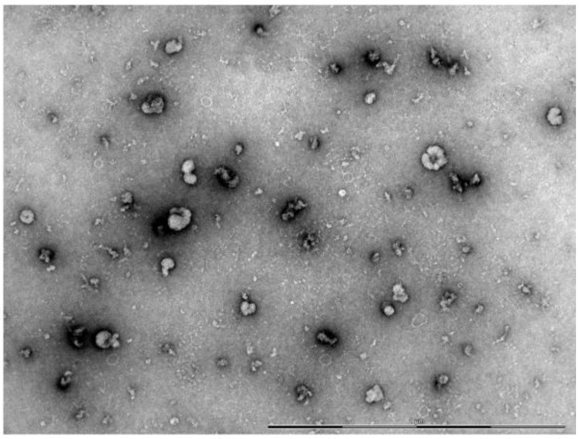

C. CD9-Tluc-CD81EVs

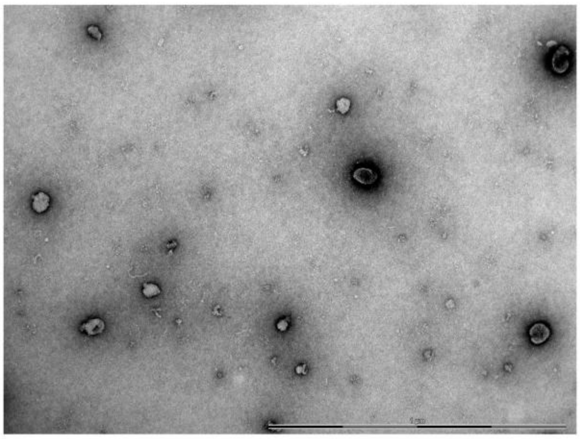

D. CD63KO-Tluc-CD81EVs

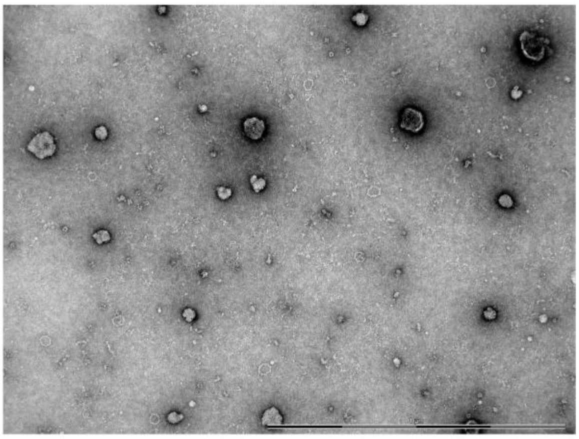

E. CD81KO-Tluc-CD63EVs

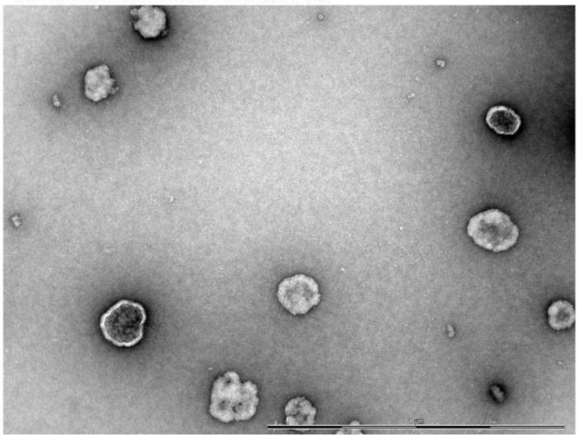

**Supplementary Figure 12**. Detection of EVs by TEM. Representative images of. A: WT-Tluc-CD81EVs. B: PanKO-Tluc-CD81EVs. C: CD9KO-Tluc-CD81EVs. D: CD63KO-Tluc-CD81EVs. E: CD81KO-Tluc-CD81EVs. Scale bar: 1 µm.

Supplementary Figure 13

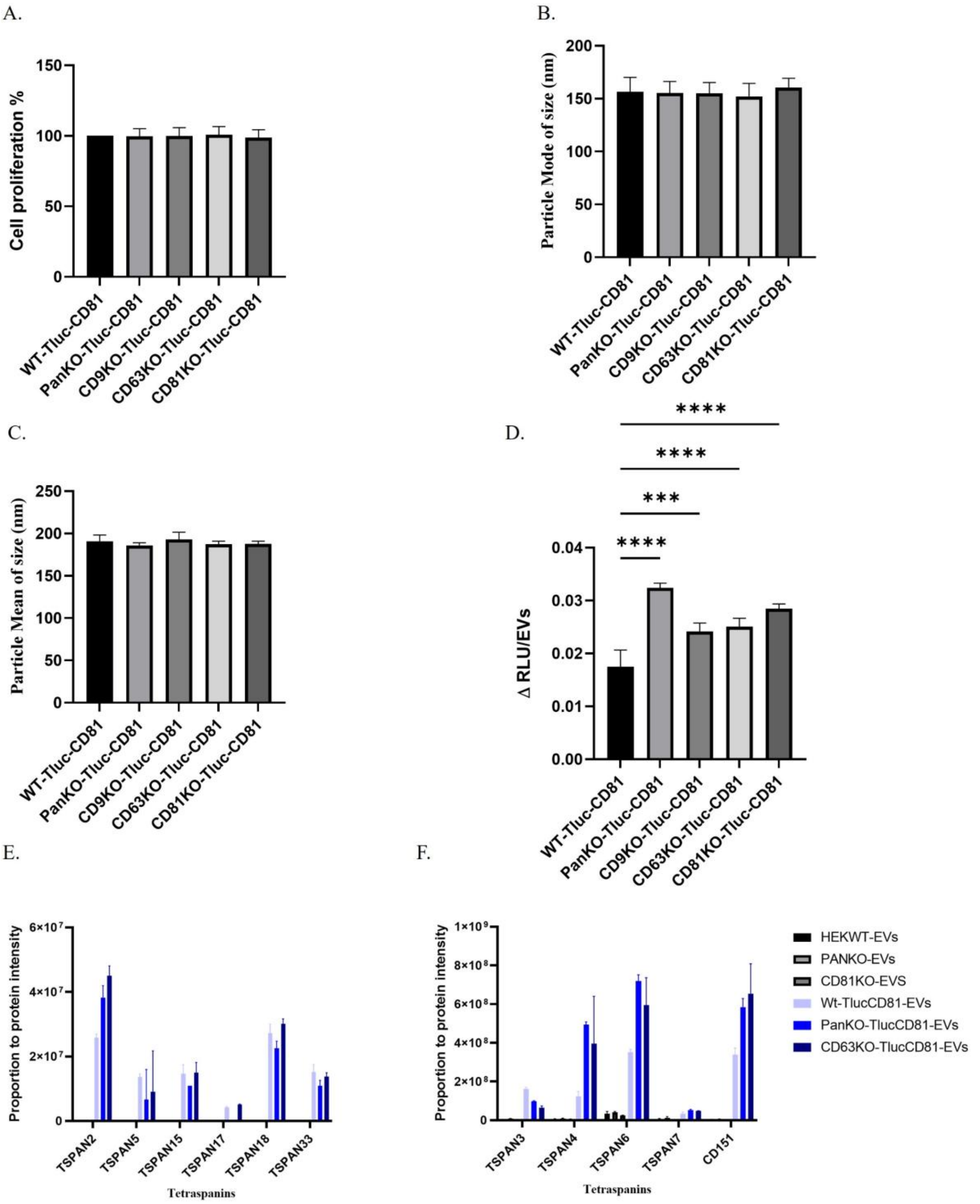


**Supplementary Figure 13**: Generation of stably expressing TlucCD81-Cerulean by lentiviruses in WT, PanKO, CD9KO, CD63KO, and CD81KO cells. A. Cell proliferation assay by performing WST-1 at absorbance of 450nm to 650nm. B. Particle mode of size in nm. C. Particle mean of size in nm. D. Relative luminescence unit (RLU) of engineered Tluc-CD81-EVs produced by KO cells and WT cells (normalized over free RLU). E and F. Expression and overexpression of tetraspanin from proteomics using MaxQuant data-driven mass spectrometry for EVs from KO and WT cells (TOTAL MS AREA/tetraspanin MS Area). The data are presented as means (±SD, n = 3-6). One-way ANOVA was used to show significance and was illustrated as follows: **** $p < 0.0001$

Supplementary Figure 14

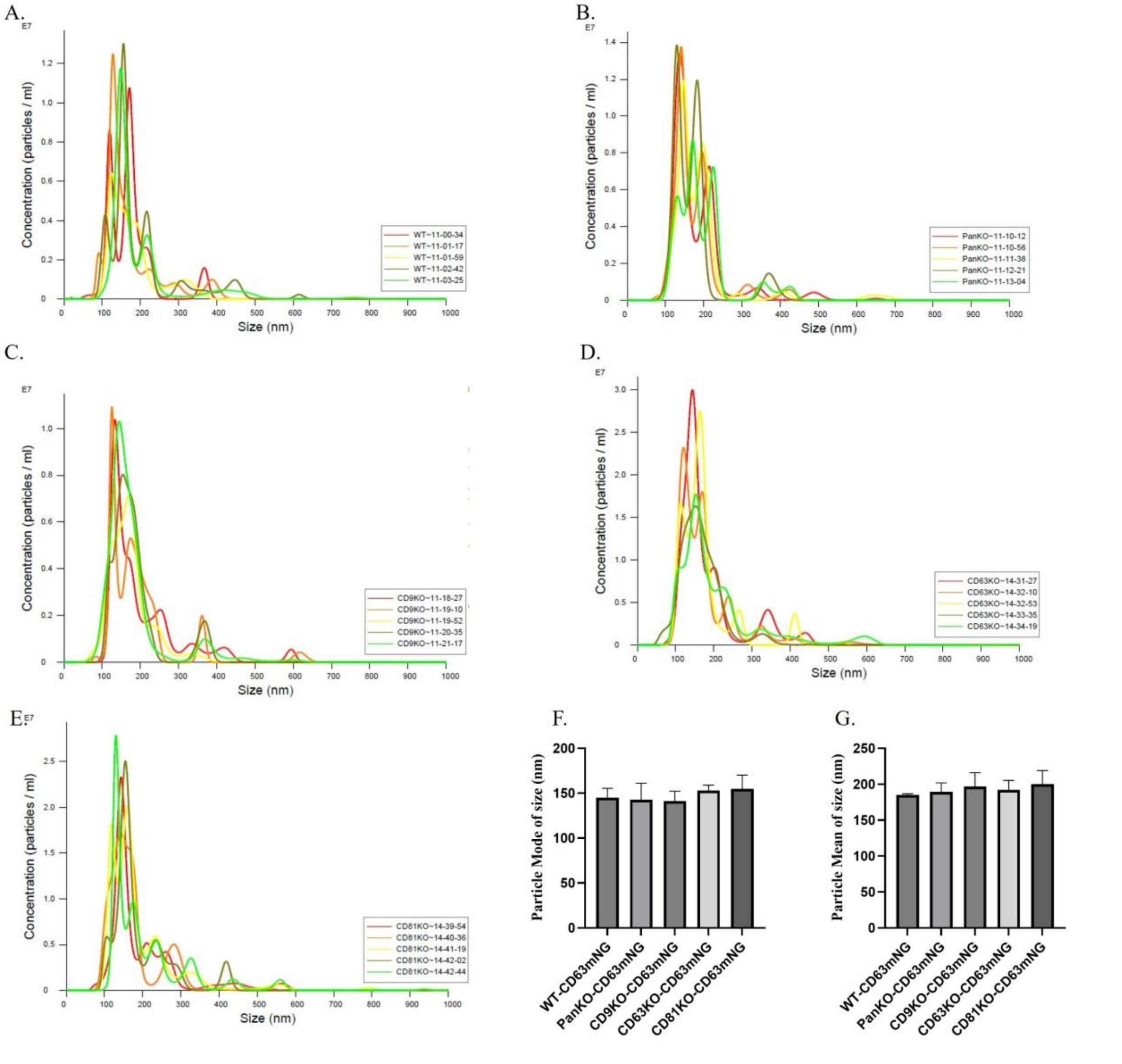


**Supplementary Figure 14**. Detection of EV size distribution using NTA. A: Size of WT-CD63mNG-EVs nm. B: Size of PanKO- CD63mNG-EVs in nm. C: Size of CD9KO-CD63mNG-EVs in nm. D: Size of CD63KO- CD63mNG-EVs in nm. E: Size of CD81KO-CD63mNG-EVs in nm. F. Particle mode of size in nm. G. Particle mean of size in nm. The data are presented as means (±SD, n = 3). NanoSight NS500 equipped with NTA 3.2 analytical software (Malvern Panalytic, UK) was used.

Supplementary Figure 15

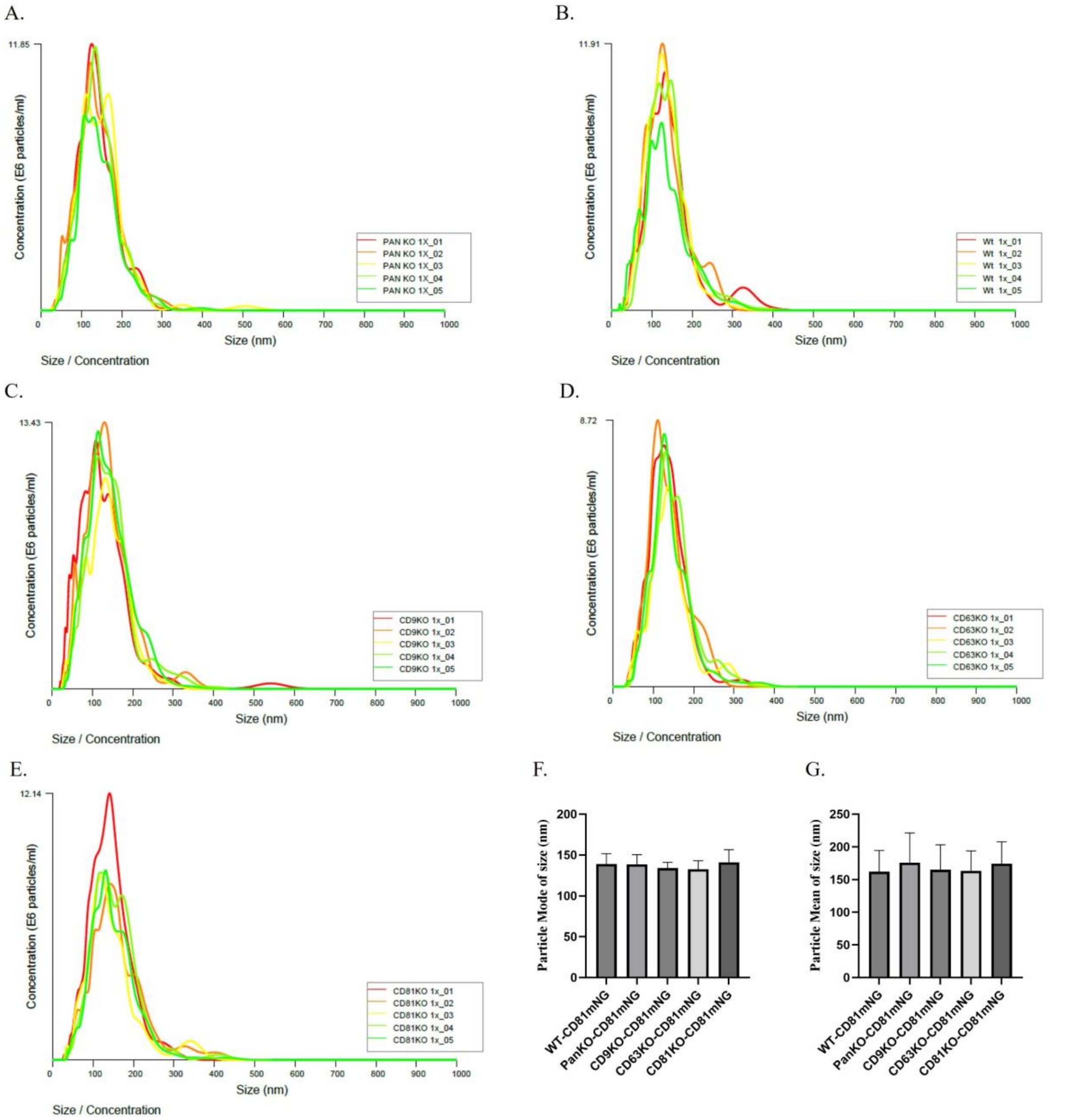


**Supplementary Figure 15**. Detection of EV size distribution using NTA. A: Size of WT-CD81mNG-EVs nm. B: Size of PanKO-CD81mNG-EVs in nm. C: Size of CD9KO-CD81mNG-EVs in nm. D: Size of CD63KO-CD81mNG-EVs in nm. E: Size of CD81KO-CD81mNG-EVs in nm. F. Particle mode of size in nm. G. Particle mean of size in nm. The data are presented as means (±SD, n = 2). NanoSight NS500 equipped with NTA 3.2 analytical software (Malvern Panalytic, UK) was used.

Supplementary Figure 16

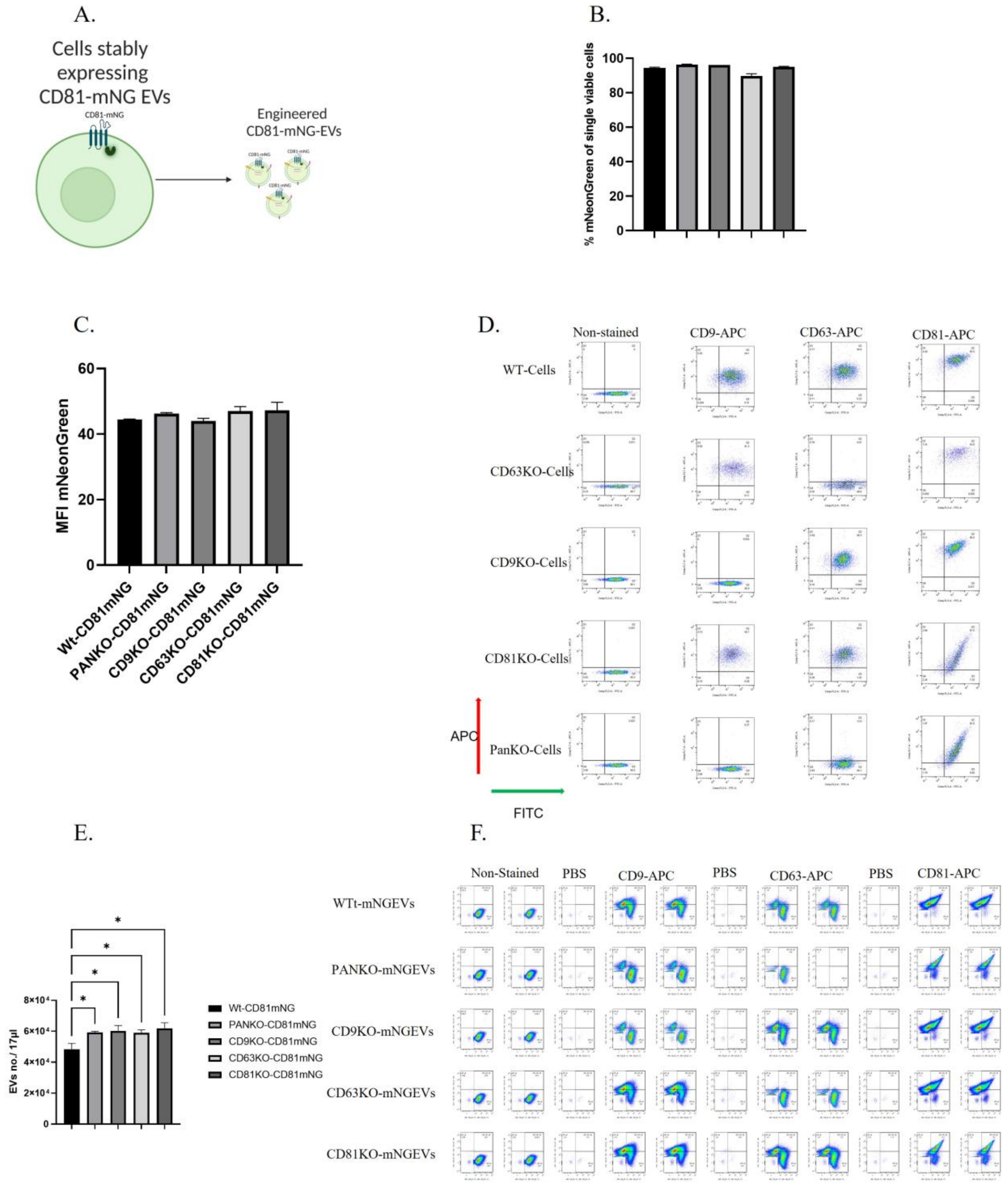


**Supplementary Figure 16**: Generation of stably expressing CD81-mNG-EVs in WT, PanKO, CD9KO, CD63KO, and CD81KO cells. A. Schematic workflow of engineered mNG-EVs by introducing CD81-mNG lentiviruses. B. Percentage of mNG positive cells after transduction using flow cytometry. C. Mean Fluorescent intensity (MFI) of the cells using flow cytometry. D. The flow cytometry plot for the cells after transduction stained with either CD9-, CD63-or CD81-conjugated antibodies. E. Number of engineered CD81-mNG EVs in KO and WT cells. F. Imaging flow cytometry plot for the mNG-EVs derived from stably expressing mNG cells. One-way ANOVA was used to show significance and was illustrated as follows: * $p < 0.05$.
